# Flowers that cool themselves: thermal ecology of summer-blooming thistles in hot Mediterranean environments

**DOI:** 10.1101/2024.11.26.625370

**Authors:** Carlos M. Herrera

## Abstract

Flower exposure to high temperature reduces the production, viability and performance of pollen, ovules and seeds, which in turn reduces individual fecundity and risks the survival of populations. Autonomous flower cooling could help to alleviate exposure of pollen and ovules to harmful temperatures, yet investigations on the thermal ecology of flowers in hot environments are still needed to evaluate the frequency, magnitude and ecological significance of floral thermoregulatory cooling. This paper reports a study on the thermal ecology of flower heads (=capitula) of 15 species of summer-blooming Asteraceae, tribe Cardueae, from hot-dry habitats in the southern Iberian Peninsula. Temperature inside (*T*_in_) and outside (*T*_out_) capitula were assessed under natural field conditions by two complementary sampling and measurement procedures, which provided information on the relationships between the two temperatures at the levels of individual capitula (“continuous recording”) and local plant populations (“instantaneous measurements”). Baselines for the *T*_in_–*T*_out_ relationship in absence of physiological activity were obtained by exposing dehydrated capitula to variable ambient temperatures. To assess whether the co-flowering capitula of summer-blooming Asteraceae collectively defined a distinct thermal layer, the vertical distribution of capitula relative to the ground was quantified. Bees visiting capitula were watched and the temperature of the air beside the visited capitulum was measured. Capitula of all species were exposed to high ambient temperatures during long periods, yet their interior was cooler than the air for most of the time, with temperature differentials (Δ*T*=*T*_in_-*T*_out_) quite often approaching, and sometimes exceeding −10°C. The relationship between *T*_in_ and *T*_out_ was best described by a composite of steep and shallow linear relationships separated by a breakpoint (Ψ, interspecific range=25°C–35°C). Capitula were weakly or not thermoregulated for *T*_out_<Ψ, but switched to intense, thermoregulated cooling when *T*_out_>Ψ. Narrow vertical distribution of capitula above ground and similar cooling responses by all species produced a refrigerated floral layer where most bees foraged at *T*_out_>Ψ and presumably visited cooled capitula. Thermoregulatory cooling of capitula can benefit plant reproduction by reducing pollen and ovule exposure to high temperature, but also the populations of insect mutualists and antagonists through habitat improvement via a refrigerated flower layer.

**OPEN RESEARCH STATEMENT:** Data, metadata and R script will be made available at figshare.

## INTRODUCTION

Plant exposure to high ambient temperature can have detrimental impacts on critical physiological, developmental and ecological processes. The photosynthetic rate of leaves (Jones 1992, Lambers et al. 2008), and the production, viability and performance of pollen, ovules and seeds (Seymour et al. 2009, Rosbakh et al. 2018, Chaturvedi et al. 2021, Arathi et al. 2023, Rosenberger et al. 2024, Tushabe and Rosbakh 2024), all decline following exposure to ambient temperatures exceeding some species-specific threshold. By combining through a variety of mechanisms, such detrimental effects of high temperature can eventually impair the fecundity of individuals and jeopardize the persistence of plant populations (Rosbakh et al. 2018, Lohani et al. 2020, Hemberger et al. 2023, Posch et al. 2024, Rosenberger et al. 2024). Since thermally-induced harmful effects on plant fitness are expected to become more frequent in the current scenario of global warming, there has been a recent upsurge of interest on the possible means whereby plants could buffer, alleviate and/or tolerate rising ambient temperatures and increasing frequency of heatwaves (Perkins-Kirkpatrick and Lewis 2020, Lorenzo et al. 2021). Most of these studies have focused on the plants’ vegetative parts and, more specifically, have addressed the possibility that the long-known ability of leaves to cool themselves to temperatures lower than the ambient (Linacre 1964, 1967, Drake et al. 1970, Pearcy et al. 1972, Ehrler 1973, Smith 1978, Upchurch and Mahan 1988) could provide a community-wide thermoregulatory mechanism allowing a “thermal escape” of leaves in the face of rising temperatures (Michaletz et al. 2016, Cook et al. 2021, Still et al. 2022, Tarvainen et al. 2022, Guo, Still et al. 2023, Manzi et al. 2024, Posch et al. 2024). Few studies, however, have explicitly examined to date how wild plants can cope with exposure of reproductive structures to high temperatures, despite the fact that the reproductive phase seems more sensitive to elevated temperatures than the vegetative one (Lohani et al. 2020, Chaturvedi et al. 2021, Tushabe and Rosbakh 2024).

Limited evidence has shown so far that autonomous flower cooling could provide an efficacious way of reducing the exposure of pollen and ovules to high ambient temperatures (Patiño and Grace 2002, Galen 2006, Rejsková et al. 2010, Karban et al. 2023, Herrera 2024a, Sherer et al. 2024). Nevertheless, investigations on the thermal ecology of flowers in hot environments are still needed to evaluate the frequency, magnitude and ecological significance of floral thermoregulatory cooling. For example, from a community viewpoint it remains to be investigated whether concurrent flower cooling by coexisting, simultaneously flowering species in hot environments could collectively produce a “refrigerated flower layer” sufficiently persistent as to be of ecological significance, e.g., by providing thermally favorable microhabitats from the perspective of pollinators (Herrera 2024a). This paper presents the results of a study on the thermal ecology of the flower heads of a group of species of Asteraceae which bloom simultaneously during the hot summer in open habitats of the southern Iberian Peninsula. The climate of the region is of a Mediterranean type, with hot-dry summers where maximum daily temperature >40°C and air relative humidity <15% are not exceptional during July-August (Capel Molina 1981, Herrera et al. 2023). Few plants bloom in southern Iberia during the climatically harsh summer (Herrera 2024a). The species of Asteraceae included in this study represent outstanding exceptions to this phenological pattern. They thus are good candidate systems to investigate floral responses to high temperatures and, more specifically, to evaluate the ability and extent of thermoregulatory cooling by species whose flowering seasons are predictably linked to hot environments. By combining population-level sampling and continuous monitoring of the internal and external thermal environments of individual flower heads, this study found substantial autonomous cooling of the interior of the flower heads in all species examined when ambient temperature exceeded a species-specific threshold. A refrigerated, multispecies floral layer arose in summer in open habitats as a consequence of the narrow vertical distribution of flower heads above the ground and their similar cooling responses to high ambient temperatures, and most bee pollinators foraged at ambient temperatures which were higher than the cooling thresholds of the visited flower heads.

## MATERIALS AND METHODS

### Study plants and field sites

Patterns of variation of temperature in the interior of flower heads (“capitula”, singular “capitulum”, hereafter), in relation to external air temperature, were studied during June– September 2020–2024 in 15 species of summer-flowering Asteraceae, tribe Cardueae, belonging to subtribes Carduinae (genera *Carduus*, *Cirsium*, *Picnomon*, *Ptilostemon*; 7 species), Centaurinae (*Carthamus*, *Centaurea*, *Mantisalca*; 4 species) and Carlininae (*Carlina*; 4 species) (Figure 1; subtribal classification follows Herrando-Moraira et al. 2019). The capitula of the species included in the study exhibited a broad variety of size, flower color and features of the bracts envelope (Figure 1). Most taxa chosen for study are widely distributed over the southern third of the Iberian Peninsula. They are often abundant in suitable habitats, where up to 7 different species can coexist locally and flower simultaneously (C. M. Herrera, unpublished data). All species typically occur in sunlit places, such as large forest clearings, roadsides, understory of sparse woodland, or extensive natural or anthropogenic disturbances, and the capitula were exposed to direct sunshine during most or all daytime. The local flowering period of each species generally encompasses 1.5–2.5 months over June–September (Blanca et al. 2011, C. M. Herrera, unpublished data). With the single exception of *Mantisalca salmantica*, all species considered here are thistles with prickly-edged leaves and densely spinescent stems and capitula (Figure 1). Most species were studied at sites in the Sierra de Cazorla area (Jaén province, 700–1600 m a.s.l.), a large mountain system in the Betic ranges that is characterized by well-preserved vegetation and high plant and habitat diversity (Gómez-Mercado 2011, Pugnaire et al. 2024). To broaden the taxonomic and microclimatic ranges represented in the study, data for four species were also collected in central Sierra Morena (Córdoba province, 760 m a.s.l.; three species) or the lowlands of the Guadalquivir River valley (Sevilla province, 25 m a.s.l.; one species), 160 and 280 km away, respectively, from the Cazorla main study area (see caption to Figure 1 for sampling locations of each species).

**FIGURE 1.**
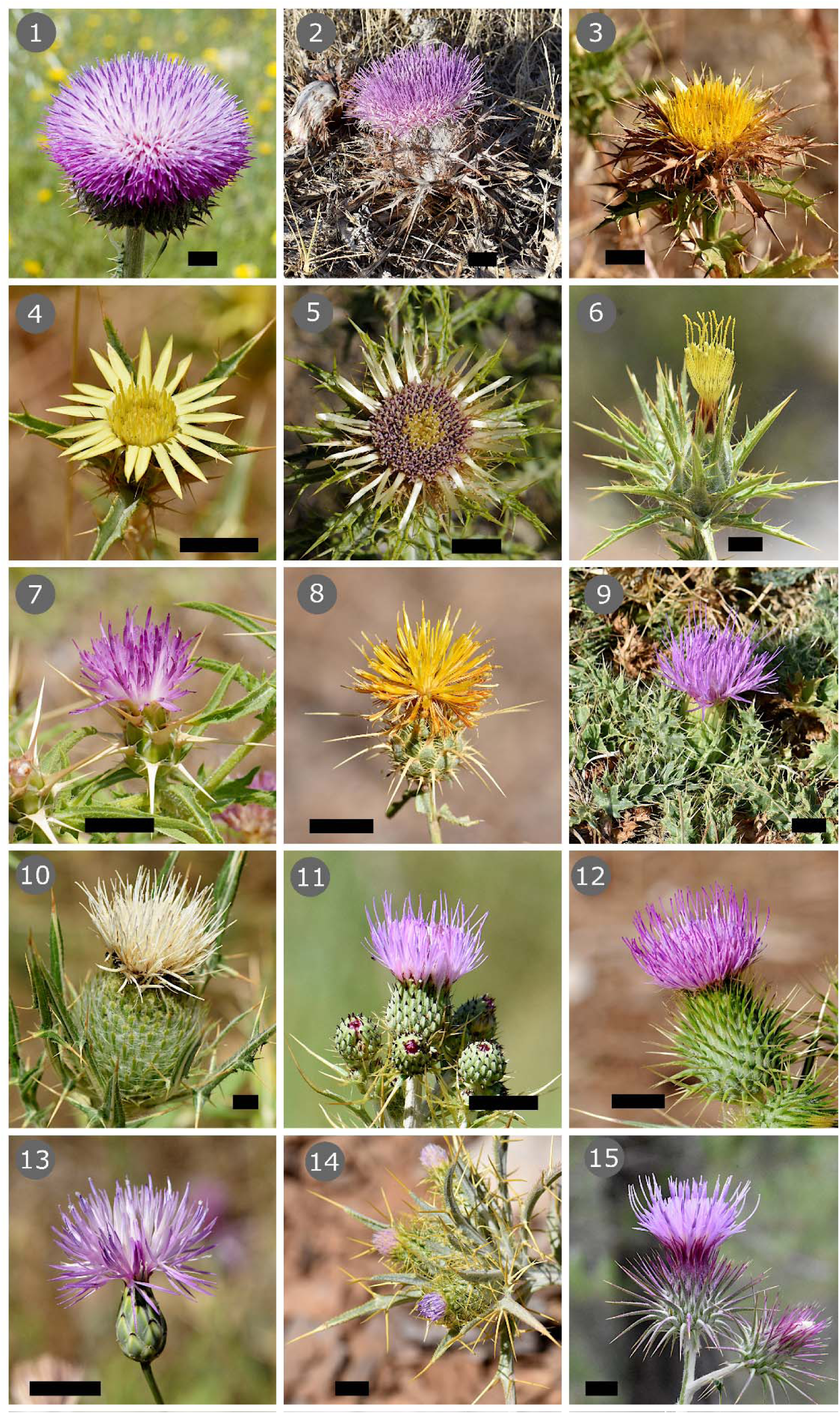
Capitula of the 15 species of summer-flowering Asteraceae, tribe Cardueae, from the southern Iberian Peninsula considered in this study. In parentheses following species names are average dry mass of individual capitula, and abbreviations for the study site(s) where was investigated each species (SC, Sierra de Cazorla; SM, Sierra Morena; LG, lowlands of Guadalquivir River valley). 1, *Carduus granatensis* (2.68 g, SC); 2, *Carlina gummifera* (6.20 g, LG); 3, *Carlina hispanica* (0.52 g, SC, SM); 4, *Carlina racemosa* (0.26 g, SC, SM); 5, *Carlina vulgaris* (0.54 g, SC); 6, *Carthamus lanatus* (0.70 g, SC); 7, *Centaurea calcitrapa* (0.18 g, SC); 8, *Centaurea ornata* (0.45 g, SM); 9, *Cirsium acaule* (0.33 g, SC); 10, *Cirsium odontolepis* (5.98 g, SC); 11, *Cirsium pyrenaicum* (0.11 g, SC); 12, *Cirsium vulgare* (1.07 g, SC); 13, *Mantisalca salmantica* (0.10 g, SC); 14, *Picnomon acarna* (0.29 g, SC); 15, *Ptilostemon hispanicus* (1.23 g, SC). Species are depicted at different scales, black bar = 10 mm.

### Field methods: natural capitula

Two complementary sampling and measurement procedures were used to assess variations of temperature inside and outside capitula under natural field conditions, which provided information on the relationships between the two temperatures at the levels of the individual capitula and the local plant populations, respectively.

In one procedure (“continuous monitoring” hereafter), paired measurements of temperatures inside individual capitula (*T*_in_ hereafter) and in the air 2 cm away (*T*_out_) were uninterruptedly recorded over periods of 2–6 days, which roughly matched in each case the functional duration of the capitulum. One fine type K thermocouple (exposed junction, 0.2 mm probe diameter, RS PRO Reference 110-4482) was inserted perpendicularly to the surface of the capitulum until it reached the receptacle. A second, similar thermocouple was attached to the capitulum and its tip remained in the air ∼2 cm away from the surface of florets (see Appendix S1, Figure S1, for examples of measurement setup). Strips of masking tape were used to attach and held the wires in place. Thermocouples were connected to a battery-powered Omega HH520 data logger thermometer. For each monitored capitulum, paired temperature measurements were continuously recorded at either 3-min (2021) or 1-min intervals (2022– 2024). A total of 57 capitula from 12 species were monitored on 91 different dates, yielding a cumulative total of 209,273 paired *T*_in_–*T*_out_ records and 178 capitulum·day of temperature data, all species combined. These data were used to assess the extent of cooling achieved by individual capitula and to examine whether regular diel rhythms existed in capitulum temperature at the individual and population levels.

The other procedure (“instantaneous measurements” hereafter) consisted of measuring *T*_in_ and *T*_out_ on a large number of randomly chosen capitula in one or more populations of each species. Measurements were taken between one hour past sunrise and noon, by inserting one type K thermocouple (similar to the ones used for continuous measurements) into the capitulum and placing the tip of another thermocouple in the air 2 cm away. The two probes were connected to a Fluke 54 IIB dual input digital thermometer/logger. Measurements were recorded at 1-s intervals for 20–25 s, and the respective averages were used as instantaneous estimates of *T*_in_ and *T*_out_ for the individual capitulum. Instantaneous measurements for a given species were taken on separate dates and populations and over a broad range of ambient temperatures. A total of 2,283 paired *T*_in_–*T*_out_ measurements from 14 species were taken on 61 different dates, all species combined. These data were used to elucidate the shape of the functional response of *T*_in_ to changes in *T*_out_, and to evaluate the ability of capitula for autonomous thermoregulatory cooling.

To assess the extent to which the capitula of summer-flowering Asteraceae actually defined a distinct multispecies layer in the habitats studied, I determined the vertical distribution of capitula in relation to the ground for nine species by measuring the height of individual capitula in a random subset of those which were sampled for instantaneous *T*_in_ and *T*_out_ measurements.

### Field methods: experimental capitula

Experiments were performed in the field to obtain species-specific baselines against which to compare the *T*_in_–*T*_out_ relationships obtained from instantaneous measurements on unmanipulated, living capitula under natural conditions. Individual capitula were severed from plants and immediately introduced into a large volume of silica gel at ambient temperature to induce a quick drying. Preliminary tests showed that in nine of the species studied the shape, color, general structure and external appearance of the capitula remained essentially unaltered following fast dehydration. In the rest of species the capitula experienced significant structural deformations and/or color changes during the drying process and were excluded from the experiments. For the capitula of the nine species chosen, baselines for the *T*_in_–*T*_out_ relationship in absence of physiological activity were obtained by pinning the dehydrated, inert capitula to styrofoam pieces, and exposing them to the sun in the field at different times over the morning under a broad range of ambient temperatures (see Appendix S1: Figure S2, for examples of experimental setup). Instantaneous measurements of *T*_in_ and *T*_out_ (*N* = 628, all species combined) were taken using the same procedure as for living capitula described in the preceding section.

### Bee visitation to capitula

Over June-September of 2022 and 2024, between early morning and noon, I watched bees that were probing capitula of six of the plant species considered here which coexisted locally at several locations in the Sierra de Cazorla where paired *T*_in_–*T*_out_ instantaneous measurements on capitula were also being taken on approximately the same dates. For each watched bee, I identified the species and measured the temperature of the air 2-5 cm away from the capitulum where it was feeding using a digital thermometer and a fast-response 0.22 mm-diameter thermocouple. I measured air temperature at the foraging sites of 388 bees on 22 different dates.

### Data analysis

Due to time, equipment or access limitations, or because of methodological restrictions (e.g., dry capitula suitable to experiments could be produced only for some species), I was unable to obtain all type of data for each of the 15 plant species considered in this study. A summary of the kind of information available for each species is shown in Appendix S1: Table 1. All the raw data used in this paper are available in Herrera (2024c).

**Table 1.**
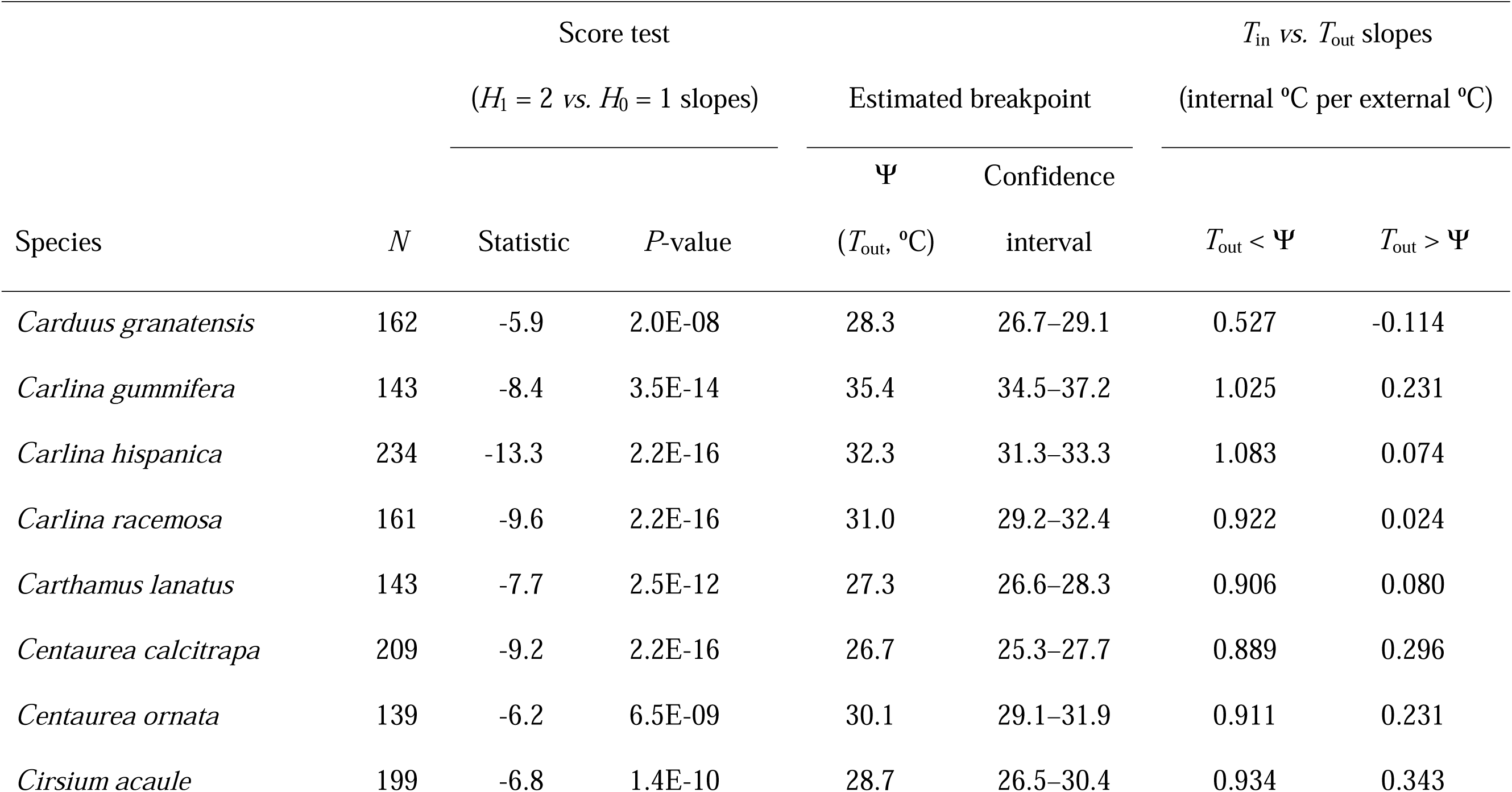
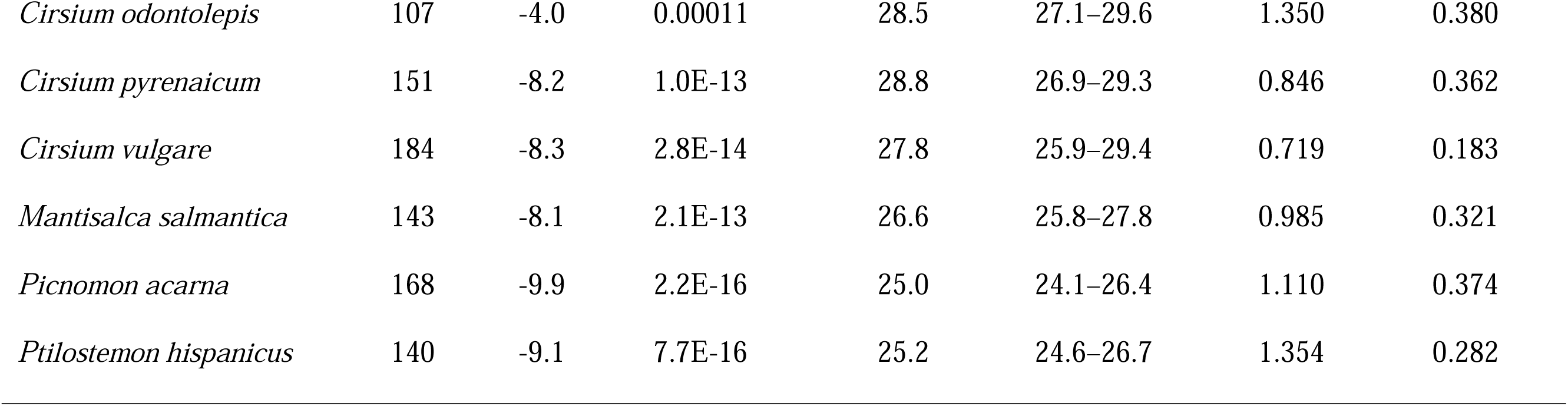
Segmented linear regression analysis of the relationship between internal capitulum temperature (*T*_in_) and temperature of the air outside (*T*_out_) for summer-flowering Asteraceae with available paired *T*_in_–*T*_out_ instantaneous measurements at the population level (*N* = number of paired measurements). See Figure 4 for plots of fitted segmented regressions.

All statistical analyses reported in this paper were carried out using the R environment (R Core Team 2024). Preliminary examination of bivariate relationships between instantaneous measurements of *T*_in_ and *T*_out_ on living capitula of all species consistently revealed non-monotonic, non-linear relationships between the two variables that could not be satisfactorily modelled using either parametric polynomial regressions or nonparametric smoothers (cubic splines). Since it was apparent that, within species, different linear relationships between *T*_in_ and *T*_out_ held for different ranges of *T*_out_, the functional relationship between the two temperatures in each set of instantaneous measurements was addressed by applying segmented regression (also sometimes named “two-phase”, “piecewise” or “broken-line” regression; Toms and Lesperance 2003, Ryan and Porth 2007). Segmented regression is a form of regression allowing multiple linear models to be fit to the data for different ranges of *x*. “Breakpoints” (Ψ) are the values of *x* at which the slope of the linear function changes, and typically they are unknown and must be estimated. Computations were performed using the segmented package, which implements a test for the existence of breakpoints (*H*_1_ = 2 slopes *vs. H*_0_ = 1 slope, over the whole *x* range sampled; function pscore.test) and provides breakpoint estimates and confidence intervals (function confint.segmented) as well as predicted values from the fitted regression (function segmented) (Muggeo 2008, 2016, 2017). The number of plant species was insufficient to perform a formal analysis to test for phylogenetic signal in the thermal parameters estimated from segmented regressions. Instead, interspecific variance in thermal parameters was partitioned into components due to subtribe, and genera nested within subtribe, using function lme in the nlme package (Pinheiro et al. 2024).

## RESULTS

### Instantaneous measurements: inert capitula

In the nine species that could be tested experimentally, the internal temperature (*T*_in_) of dehydrated, inert capitula exposed to natural sunny conditions in the field increased linearly with the temperature of the surrounding air (*T*_out_), and variation in *T*_out_ accounted for nearly all variance in *T*_in_ (91–98%, Figure 2). The linear slopes of the *T*_in_–*T*_out_ relationships were close to unity in all cases, and the capitulum-air thermal gradient (Δ*T* = *T*_in_-*T*_out_) predicted from the regressions was always positive and tended to remain between +5°C and +10°C over the whole range of ambient temperatures under which the tests were conducted (Figure 2). Under natural field conditions, therefore, inert capitula consistently built up a substantial thermal excess over their immediate environment which tended to remain fairly constant across the broad range of ambient temperatures sampled.

**FIGURE 2.**
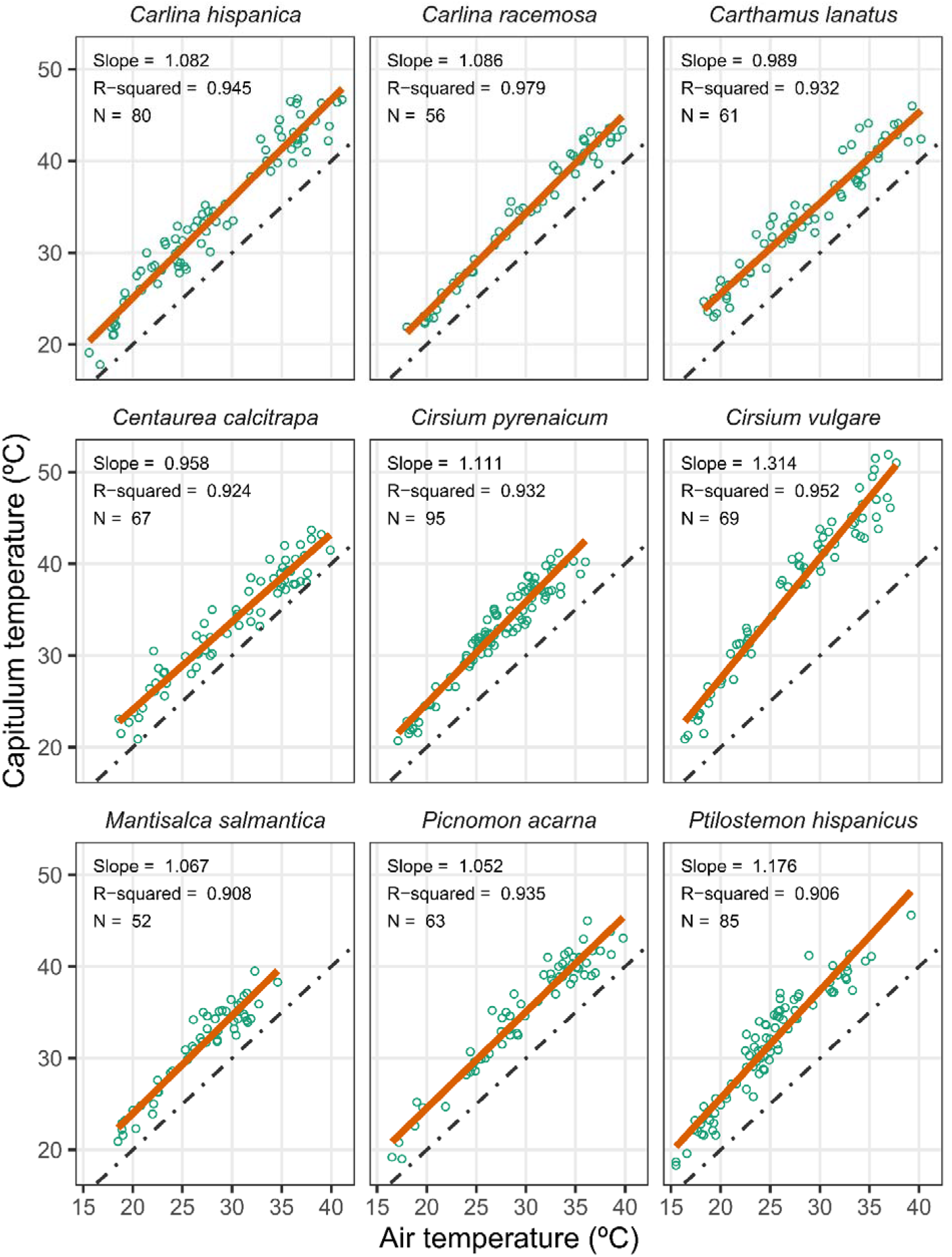
Relationship between the temperature inside the capitulum and the temperature of the surrounding air (grey dots) in samples of inert, experimentally dried capitula exposed in the field to natural variations in the thermal environment. Red lines are ordinary least-squares linear regressions fitted to the data (insets show summary statistics). Dot-dash lines represent the *y* = *x* isothermal condition.

### Instantaneous measurements: living capitula

The temperature inside living, unmanipulated capitula under natural field conditions (*T*_in_) increased with the temperature of the surrounding air (*T*_out_) in all species, but the shape of the relationship departed in every case from a simple linear, monotonously increasing relationship of the sort exhibited by the dehydrated, inert capitula (Figure 3). Segmented regression analyses rejected for all species the null hypothesis of a single slope over the whole range of *T*_out_ in favor of the alternative hypothesis of two distinct slopes separated by a breakpoint (Table 1, Figure 3). Estimated breakpoints varied widely among species (range Ψ = 25.0–35.4°C), with the highest values consistently occurring in species of *Carlina* (Ψ = 31.0–35.4°), followed by species of *Cirsium* (Ψ = 27.8–28.8°C) (Table 1).

**FIGURE 3.**
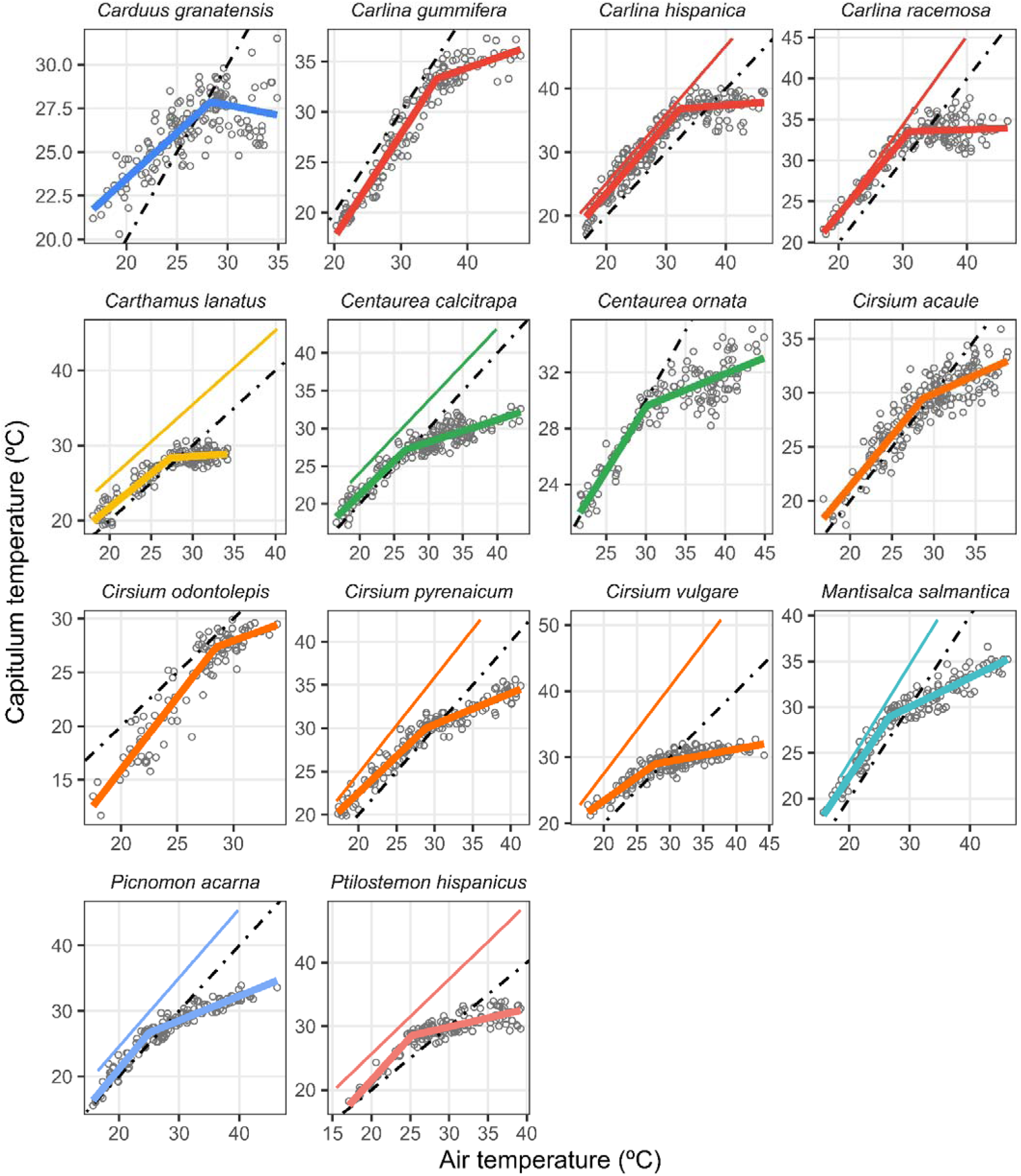
Bivariate relationships between the temperature inside capitula and the temperature of the surrounding air. Thick and thin colored lines are regressions fitted to field measurements on living (grey dots, segmented regressions, see Table 2 for details) and inert dried capitula (ordinary linear regressions shown in Figure 2), respectively, color-coded according to plant genus. For each species, paired temperature measurements were taken throughout the morning (06:00–12:00 h UTC) on capitula from different dates and populations. Dot-dash lines represent the *y* = *x* isothermal condition.

**Table 2.**
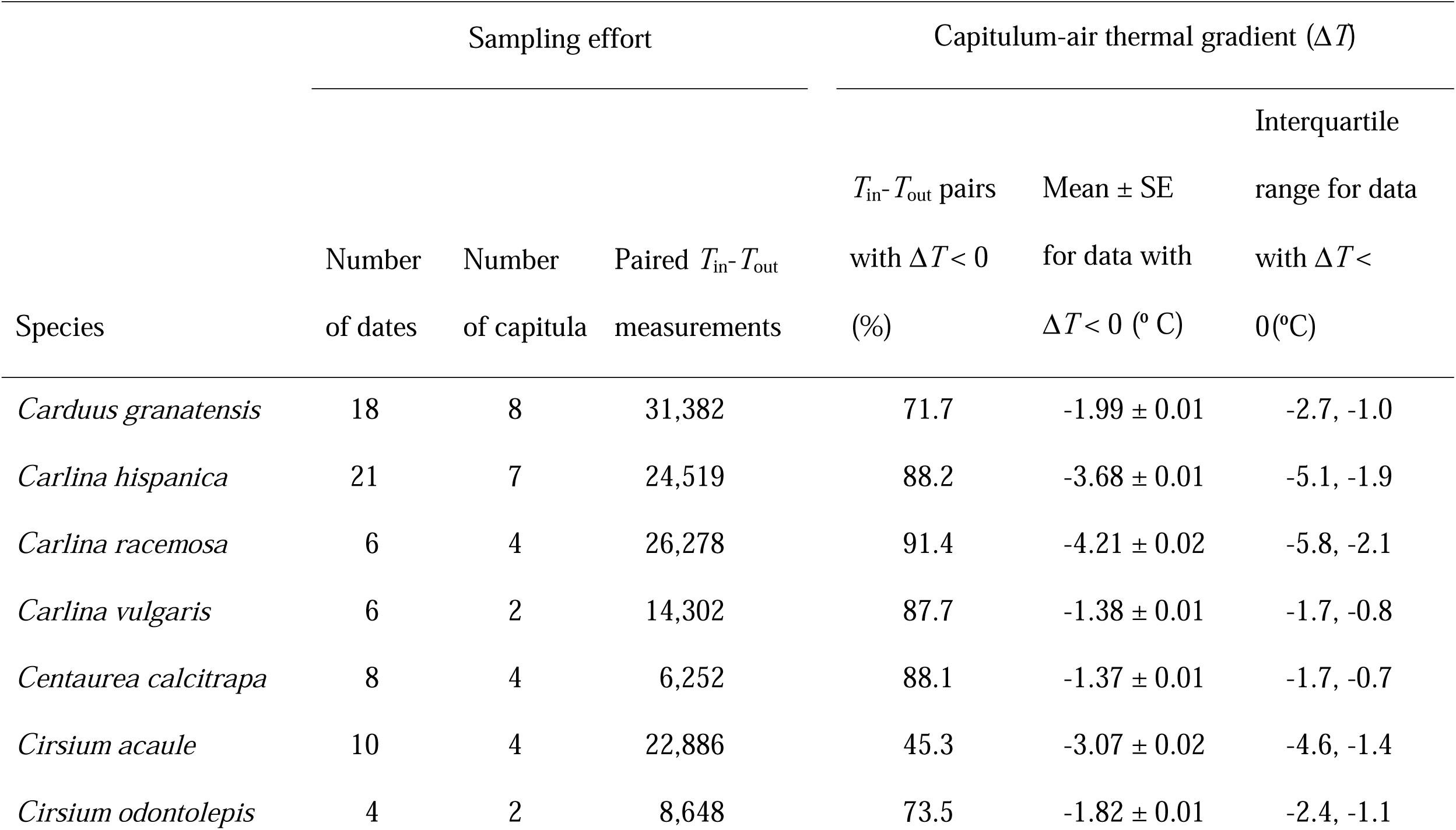
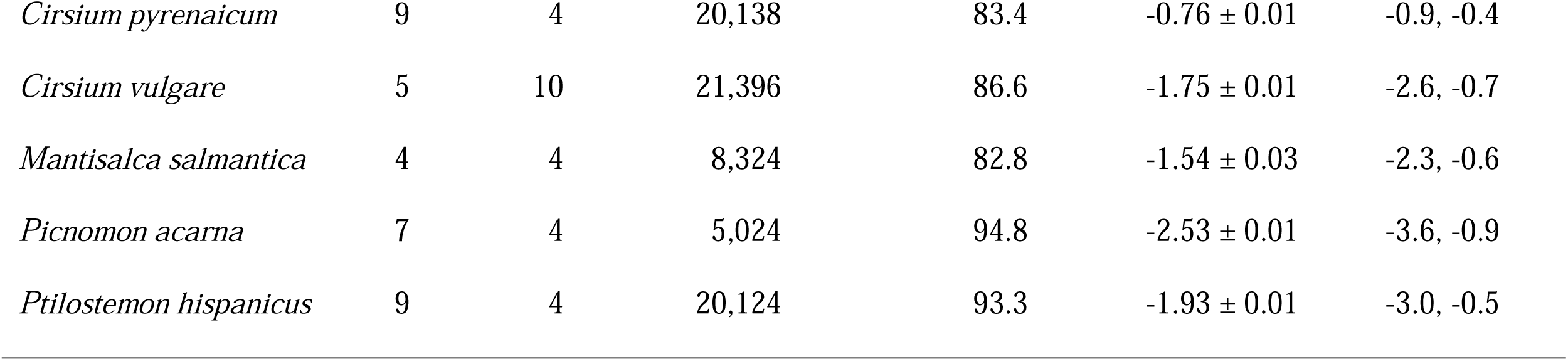
Species for which temperatures inside (*T*_in_) and outside (*T*_out_) individual capitula were continuously monitored over one or more days, sampling effort, and summary statistics of the distributions of capitulum-air thermal gradient (Δ*T* = *T*_in_ – *T*_out_) for data points with Δ*T* < 0.

At ambient temperatures lower than breaking points the internal temperature of living capitula in the field tended to fall somewhere between the two baselines represented by the *T*_in_– *T*_out_ relationship for dehydrated capitula, on the upper side, and the isothermic line *y* = *x*, on the lower one, while it fell well below these two reference lines when *T*_out_ exceeded Ψ (Figure 3). At the highest air temperatures sampled (35–48°C), temperatures within the living capitula mostly were 5–14°C lower than air temperature, and up to 15–20°C lower than those reached by the dehydrated, inert capitula (Figure 3).

Irrespective of interspecific differences in Ψ values, the overall shape of the segmented functional relationships linking *T*_in_ and *T*_out_ was remarkably similar for all species: steep slopes close to unity for *T*_out_ < Ψ and a shift to distinctly shallower slopes for *T*_out_ > Ψ (Table 1, Figure 3). Consequently, at ambient temperatures under Ψ the internal temperature of capitula generally tended to run parallel and remain fairly close to an isothermal relationship with the ambient, while an internal thermal deficit arose at *T*_out_ beyond Ψ which increased steadily as ambient temperature increased. This shift in the *T*_in_–*T*_out_ relationship denoted a significant decoupling of *T*_in_ in relation to *T*_out_ when Ψ was exceeded, which was shared by all species to a greater or lesser extent. Such decoupling was remarkable in species of *Carlina* and *Carthamus*, in which the temperature inside the capitulum became independent of ambient temperature when the latter exceeded Ψ (slopes close to zero for *T*_out_ > Ψ; Table 1 and Figure 3).

Interspecific variance in Ψ was largely accounted for variation among subtribes (71.3% of total), while the contribution of the differences among genera within tribes was comparatively negligible (6.7%) (Appendix S2: Table S1).

### Continuous monitoring

Continuous temperature measurements over most or all the functional life of individual capitula revealed that the capitulum-air thermal gradient (Δ*T*) was predominantly negative in every species except *Cirsium acaule* (Table 2). The proportion of measurements with Δ*T*<0 ranged between 72–95% among species, which denotes that the interior of individual capitula was consistently cooler than the surrounding air for most of their lives. Considering only those data with Δ*T* < 0,the capitulum-air thermal gradients averaged between −4.2°C (*Carlina racemosa*) and −0.8°C (*Cirsium pyrenaicum*) (Table 2).

Both *T*_in_ and Δ*T* followed regular diel rhythms in the individual capitula of all species, as illustrated by the time series for representative capitula depicted in Figure 4. With only minor variations among species, the prevailing pattern involved *T*_in_ being roughly similar to *T*_out_ (Δ*T*∼0) from early morning to shortly before noon, as both temperatures tended to increase in unison over that period. Subsequently *T*_in_ and *T*_out_ followed increasingly divergent courses, with substantial negative capitulum-air gradients building up which persisted for the rest of daytime (*Centaurea*, *Cirsium*, *Picnomon*), until around midnight (*Carduus*, *Ptilostemon*) or, in the most extreme cases (*Carlina*), up to the following sunrise. In some taxa, the building-up of negative Δ*T* started immediately after a short spike of Δ*T*>0 (see, e.g., series for *Carduus granatensis*, *Carlina racemosa*, *Carlina vulgaris* and *Cirsium vulgare* in Figure 4). In species whose capitula lasted for several days the absolute value of the daily peaks in negative Δ*T* tended to decline with capitulum age (e.g., *Carduus granatensis*, *Carlina racemosa*, *Centaurea calcitrapa*; Figure 4).

**FIGURE 4.**
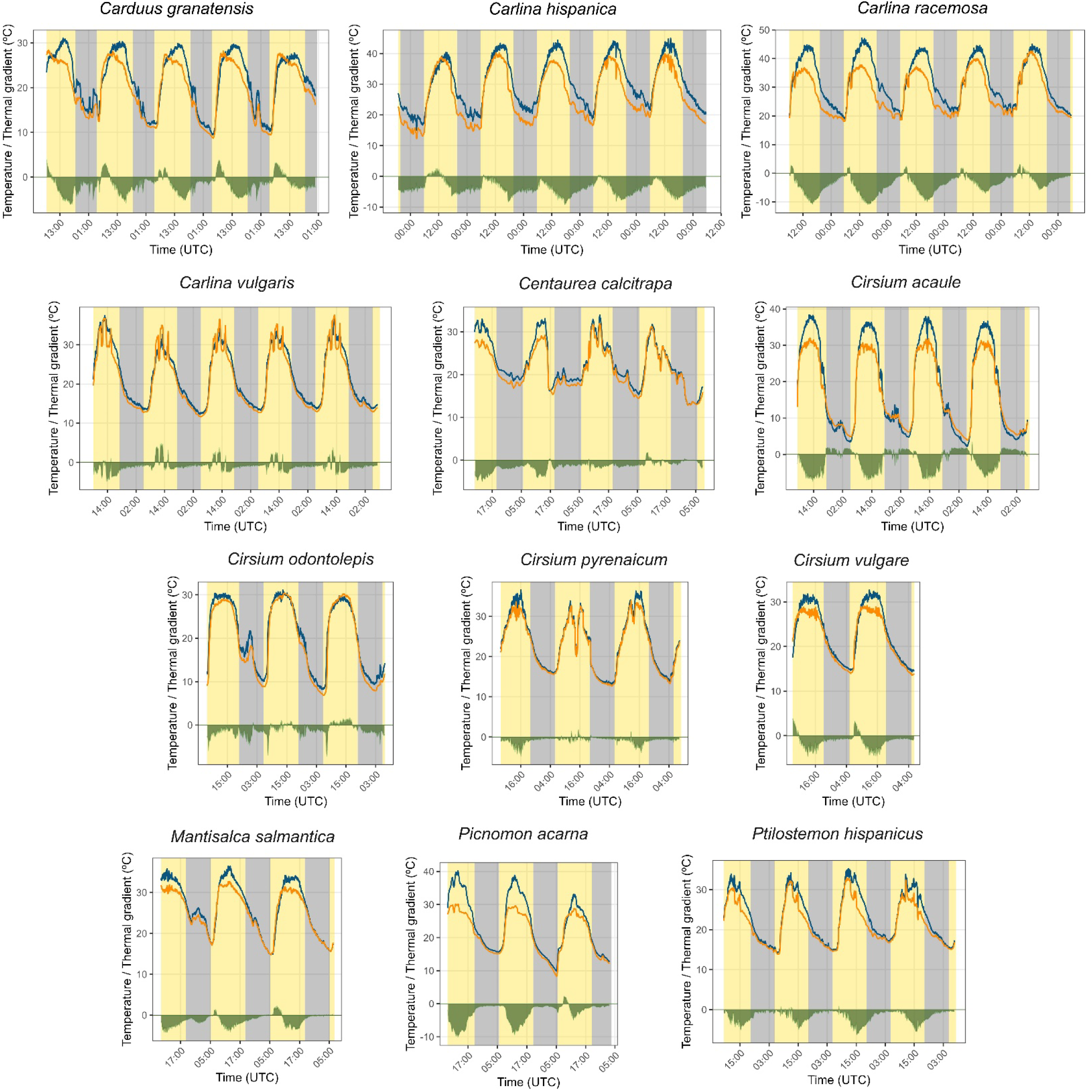
Variation of temperature inside representative capitula (one capitulum per species; *T*_in_, orange lines;), the air outside (*T*_out_, blue lines), and capitulum-air thermal gradient (Δ*T* = *T*_in_ – *T*_out_, green area at the bottom of each plot, with bottom- and top-facing areas around *y* = 0 reflecting negative and positive gradients, respectively). Original measurements were smoothed for the graphs using a moving average procedure with 10-min sliding window. Differences between species in the length of measurement period mosty reflect interspecific differences in the duration of capitula. Daytime and nighttime periods are denoted by yellow and grey vertical bands, respectively.

### Diel and spatial patterns at community level

The combination of the regular diel rhythms in Δ*T* which took place at the level of individual capitula led to distinct diel rhythms in Δ*T* at the plant population level in the majority of the species studied (Figure 5). With minor departures (e.g., *Cirsium odontolepis*) Δ*T* was, on average, large and consistently negative in the flower populations from around mid morning until late afternoon (Figure 5). Since the capitula of most species are spatially concentrated within 50 cm of the ground (Figura 6), the combined diel rhythms in Δ*T* created a multispecific layer of refrigerated capitula near the ground which lasted for a substantial part of daytime and, in some species, extended also into nighttime.

**FIGURE 5.**
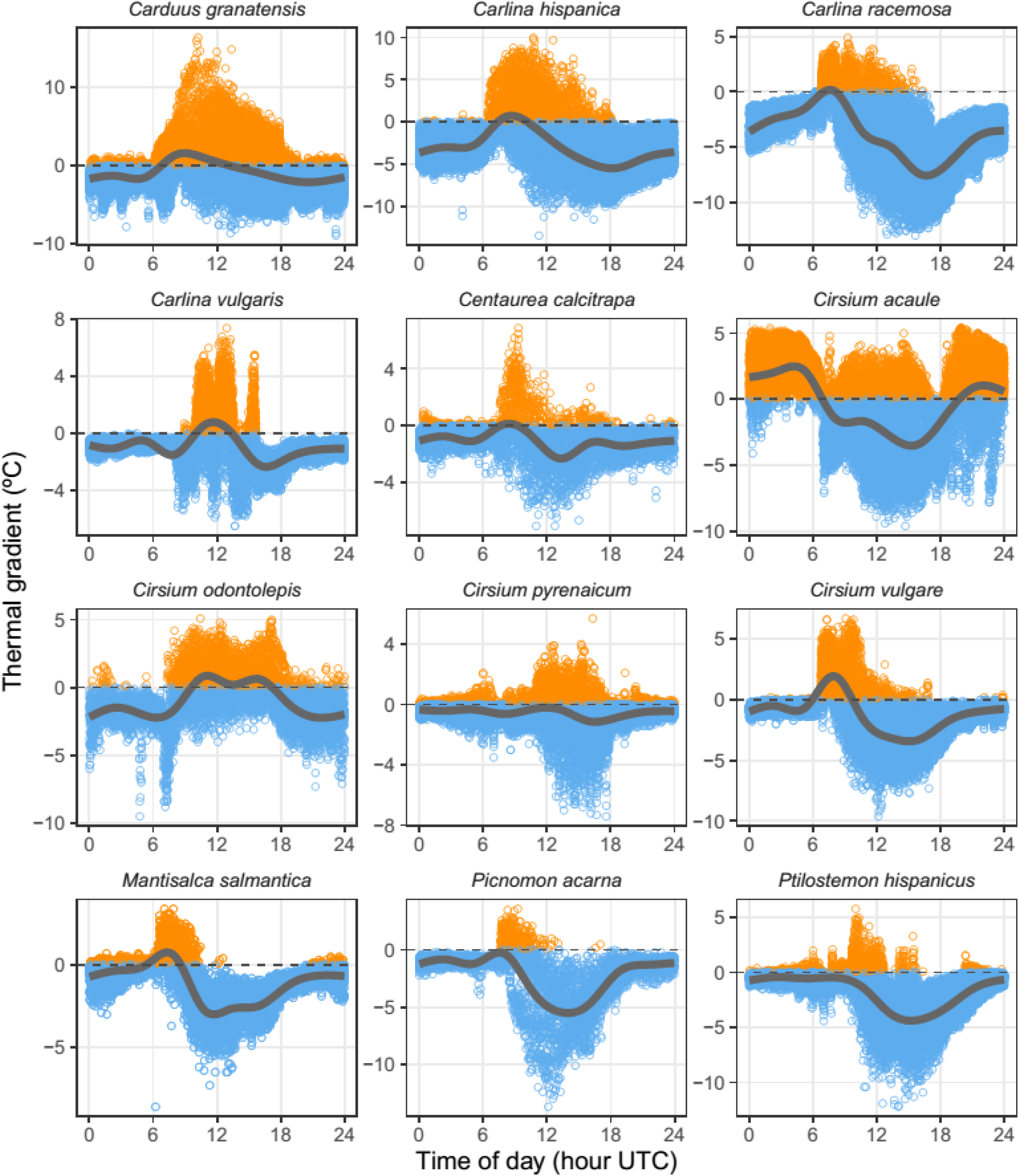
Daily variation in capitulum-air thermal gradient (Δ*T* = *T*_in_ - *T*_out_; positive and negative gradients are shown as orange and blue symbols, respectively) for the whole set of individual capitula that were continuously monitored over most or all their lifetimes. Each symbol correspond to a pair of *T*_in_ and *T*_out_ measurements (*N* = 209,273, all species combined). Data from all dates and individual capitula are combined in the plot for a given species (see Table 2 for sample sizes). The curves are cubic splines fitted to the data.

**FIGURE 6.**
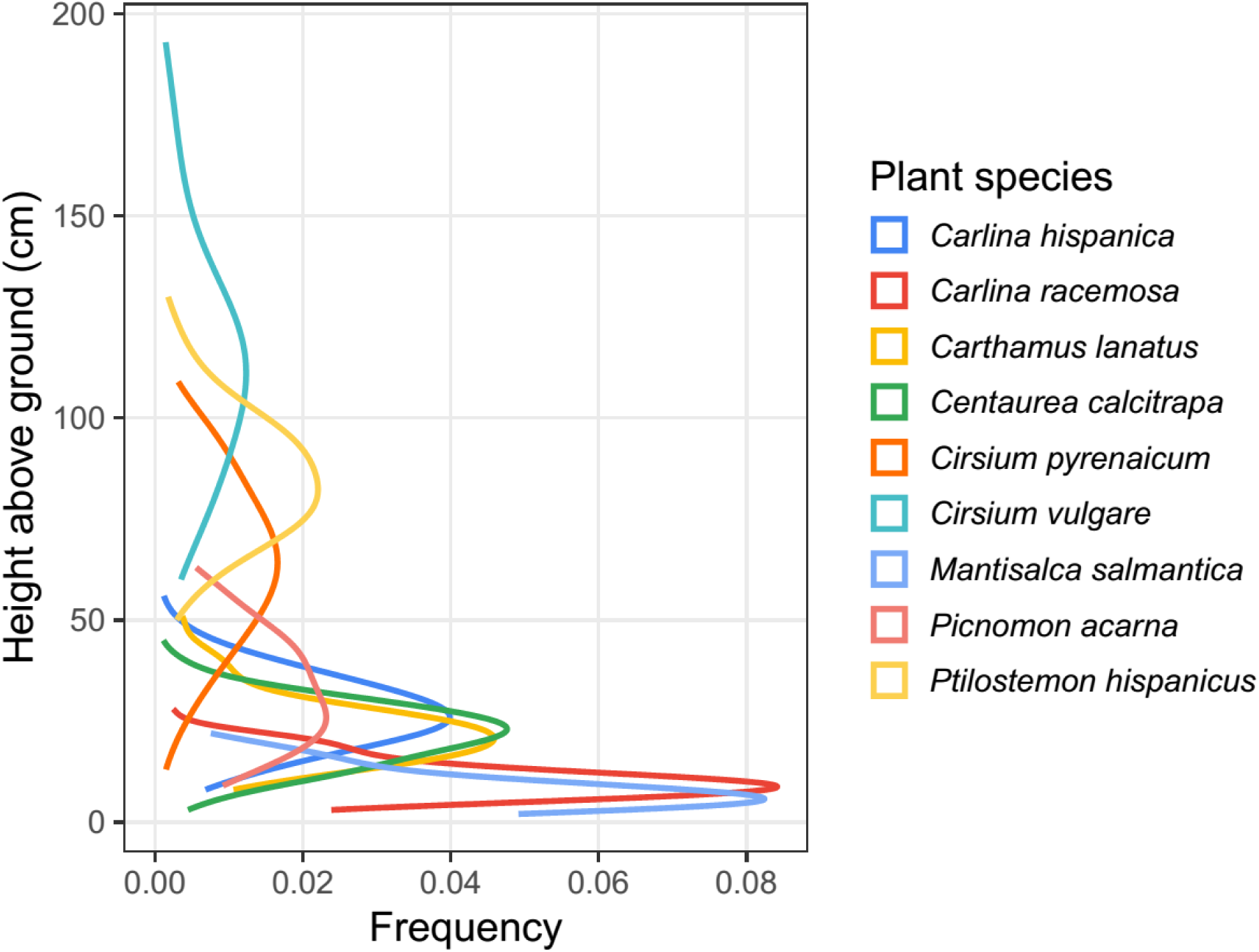
Frequency distributions of capitula height above the ground for nine of the species studied. In *Carlina gummifera* and *Cirsium acaule* all the capitula occur within 5 cm from the ground (Figure 1), and the narrow height distributions for these two species have been omitted from the graph for clarity. Curves depict the fitted kernel estimates of data point density for each species.

### Bee visitation

A total of 33 bee species were recorded visiting capitula of the six locally coexisting species chosen for field observations. The family Megachilidae was the most important numerically, contributing 57.2% of individuals and 48.5% of species, followed by Apidae (33.2% and 33.0% of individuals and species, respectively). Observations were very unevenly distributed among plant and bee species, and most bee species were rarely recorded (Appendix S2: Table S2), hence data for all bees and plants have been combined into a single sample for the analysis.

Bees were observed foraging under a broad range of air temperatures (range = 16–37°C, median = 29°C, interquartile range = 27–32°C) (Figure 7). Except for one plant species, air temperature beside the visited capitula was in the majority of instances higher than the thermal breakpoint (Ψ) for the plant species involved (Figure 7). In other words, a substantial proportion of the capitula visited by bees were expected to have already entered the active cooling stage (i.e., *T*_out_ > Ψ): 79.0% (*Mantisalca salmantica*), 77% (*Centaurea calcitrapa*), 68.0% (*Cirsium vulgare*), 59.0% (C*irsium odontolepis*), 55% (*Cirsium pyrenaicum*), and 16% (*Carlina hipanica*).

**FIGURE 7.**
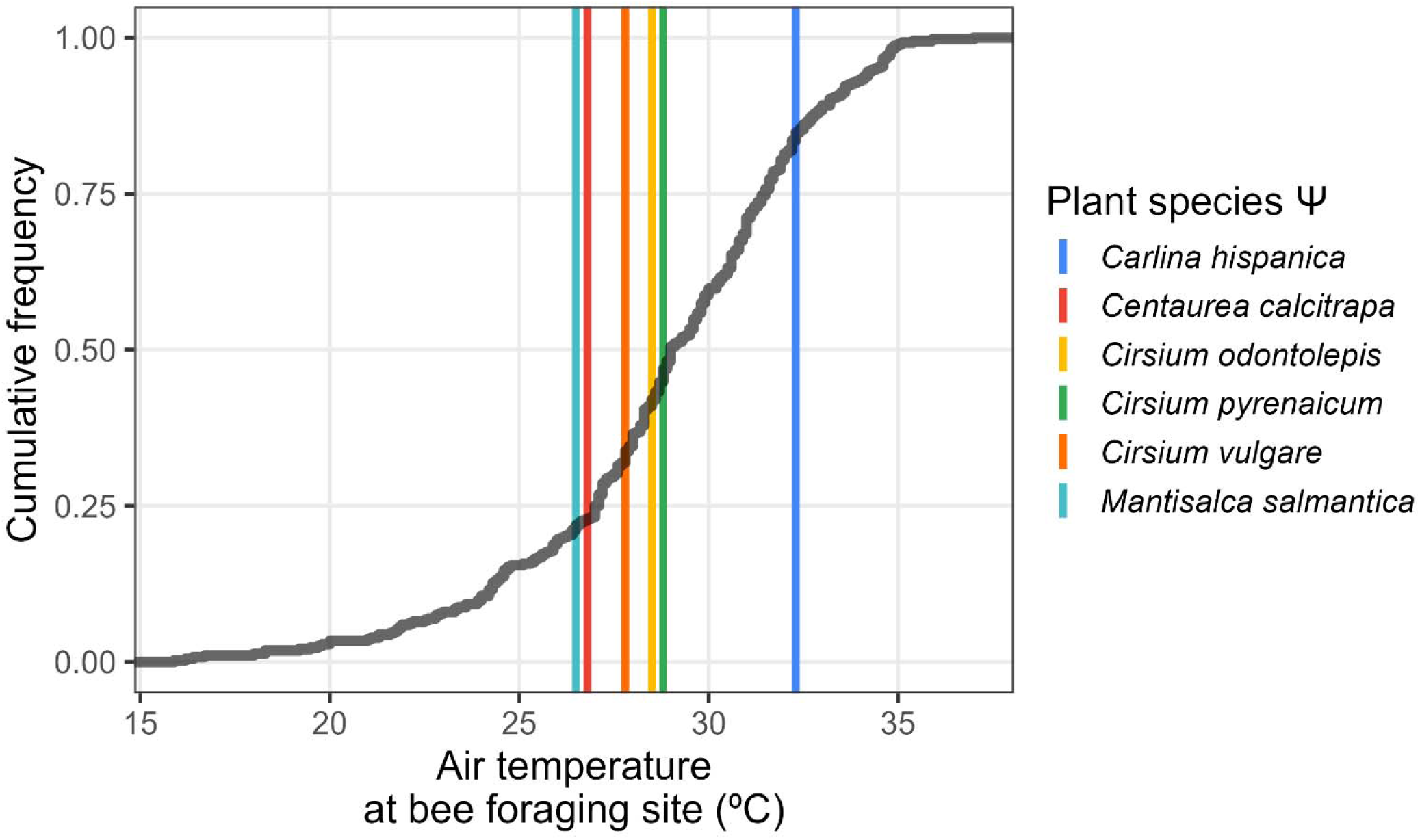
Cumulative frequency distribution of air temperature at the foraging sites of bees watched probing the capitula of six species considered in this study (grey line, *N* = 388 observations, all plant and bee species combined, see Appendix S2, Table S2 for details). Colored vertical lines mark the air temperature breakpoints (Ψ) for each plant species, as obtained from the segmented regressions of capitulum temperature (*T*_in_) against ambient temperature (*T*_out_) (Table 1).

**FIGURE 8.**
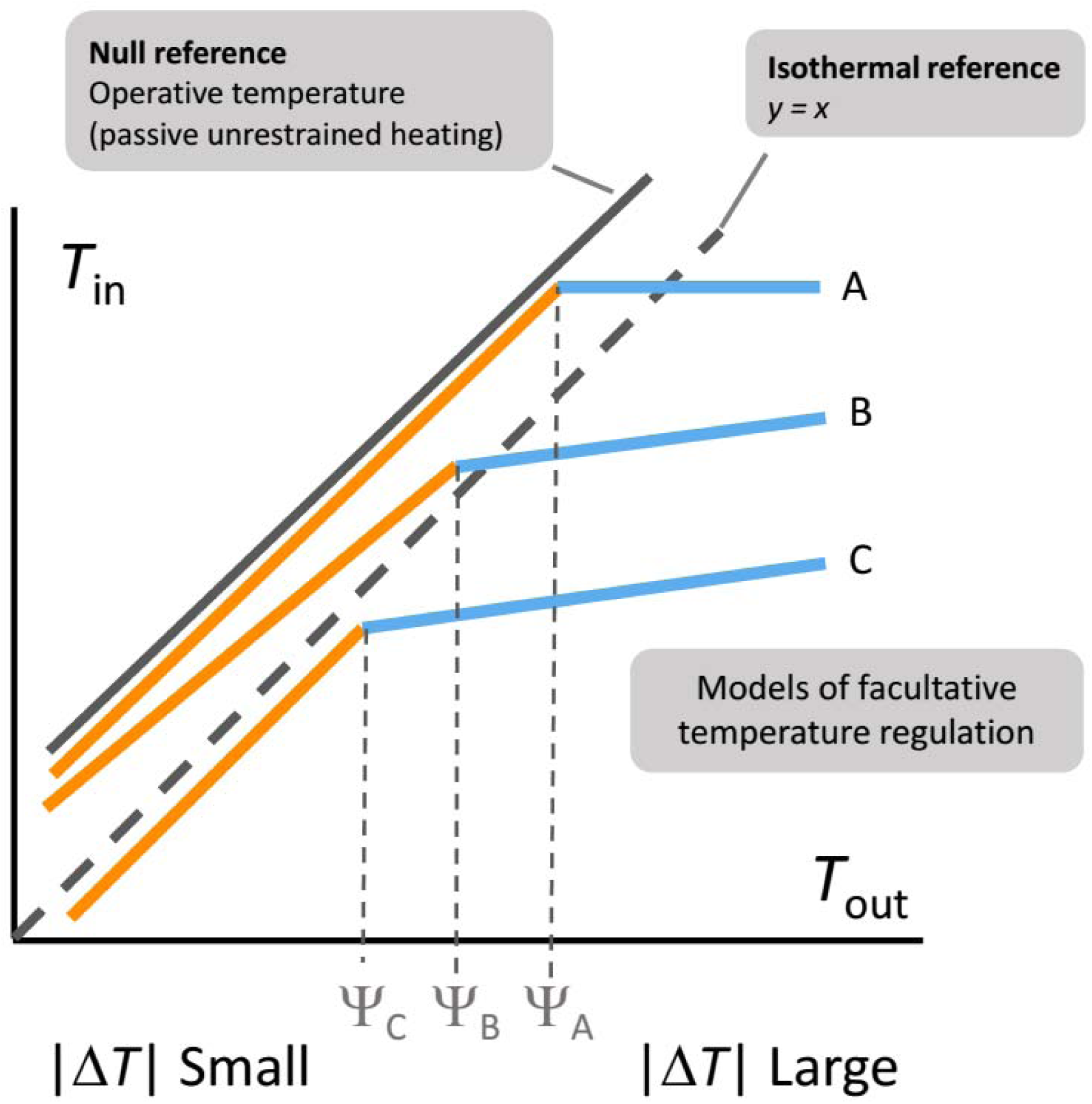
An expanded formulation of the single-slope, classical reference model of Huey and Slatkin (1976) linking organismal and ambient temperature to accomodate the two-slopes, facultative thermoregulation patterns exhibited by the capitula of Asteraceae investigated here (Figures 3). In this expanded framework, interspecific differences in thermoregulatory capacity will depend on a combination of the position of regression lines relative to the null-model and isothermal *T*_in_–*T*_out_ reference lines, the comparative values of the two regression slopes involved, and the position of break points (Ψ) along the gradient of environmental temperature. Δ*T* is the capitulum-air thermal gradient, *T*_in_ - *T*_out_. See Figure 3 for the actual relationships exhibited by the species considered in this study. Examples for the three hypothesized patterns of facultative cooling patterns, taken from Figure 3: A, *Carlina hispanica*, *C. racemosa*; B, *Cirsium pyrenaicum*, *C. vulgare*; and C, *Carlina gummifera*, *Cirsium odontolepis*.

## DISCUSSION

### Cooling

Instantaneous measurements and continuous monitoring revealed that the capitula of all the species of summer-blooming Asteraceae studied were regularly exposed in the field to long periods of high ambient temperatures (quite often >30°C, at times >40°C), yet the interior of the capitula was cooler than the surrounding air for most of their lives. The capitulum-air thermal gradient exhibited diel rhythms whose shapes varied relatively little among species and were remarkably constant over consecutive days for a given capitulum. As a general rule, the magnitude of negative capitulum-air gradients peaked around the hottest period of daytime, when they often approached or even exceeded −10°C. As far as I know, these are the largest negative temperature differentials ever reported for flowers, as well as one of the first documented examples of diel rhythms in floral temperature differentials. Comparable results were advanced by Herrera (2024a) for *Carlina corymbosa* (a junior synonym of *C. hispanica*, the name used here following Blanca et al. 2011). The few studies that have so far examined documented the ability of flowers to cool themselves either failed to document negative flower-air gradients (Patiño and Grace 2002, Galen 2006) or, when such negative gradients were found, the magnitude was much smaller than that found here for the capitula of summer-flowering Asteraceae (e.g., −2.7°C in *Galanthus nivalis*, Rejsková et al. 2010; −1.9°C in *Hexastylis arifolia*, Sherer et al. 2024). Taken together, however, these results illustrate that floral cooling can be more frequent in nature than its usual neglect in the literature on floral thermal ecology would suggest, as it was already anticipated by van der Kooi et al. (2019). These authors emphasized that the prevalence of studies on floral warming in the literature mostly reflects geographical biases in the field of floral thermal ecology rather than a biological reality, because most work on floral thermal ecology has been conducted in alpine, arctic and mid-latitude plants. Results of the present study support their view, and also prompt the prediction that future investigations on the floral thermal ecology of plants inhabiting hot-dry environments akin to those prevailing during Mediterranean summers should reveal more examples of significant autonomous cooling.

One important result of this study was the finding that the functional relationship linking internal temperature of the capitulum with ambient temperature was not a simple, monotonously linear or quasi-linear one, as frequently assumed in research on thermal ecology of leaves and flowers (Michaletz et al. 2015, 2016, Sherer et al. 2024). In all species studied here the functional response was best described by a composite of one steep and one shallow linear response separated by a (statistically estimated) breakpoint (Ψ), or “thermal threshold”. In practical terms, this finding suggests that substantial floral cooling (i.e., that occurring when *T*_out_ > Ψ) can remain unnoticed for a given plant species if the range of ambient temperatures sampled either is not broad enough to include the species-specific thermal threshold Ψ, and/or the statistical power to reject the null hypothesis of a single slope by means of segmented regression is too small because of insufficient sample size (Muggeo 2016). Furthermore, since the species-specific Ψ values above which cooling occurred spanned over ∼10°C, interspecific differences in cooling ability could sometimes be a spurious consequence of sampling thermal ranges that include the thresholds for some species but not for others.

Two-slopes functional relationships between plant and ambient temperatures of the sort documented here for capitula of Asteraceae are often discernible in, or can be inferred from, *T*_in_– *T*_out_ plots from early studies on leaf thermal ecology (e.g., Fig. 1 in Linacre 1964; Fig. 5 in Linacre 1967; Fig. 1 in Palmer 1967; Fig. 42 in Pearcy et al. 1972; Fig. 3 in Upchurch and Mahan 1988). Furthermore, the existence of underlying thermal “switching points” beyond which leaf cooling occurs, roughly equivalent to the Ψ thresholds analytically estimated here for capitula, was already documented nearly 40 years ago (Althawadi and Grace 1986, Upchurch and Mahan 1988). These parallelisms suggest that the underlying mechanism(s) of plant response to high temperature is probably similar in leaves and flowers. In the case of leaves, it seems well established that high temperatures activate stomatal opening through a photosynthesis-uncoupled signalling pathway, which enhances transpiration and leads to evaporative cooling (Drake et al. 1970, Althawadi and Grace 1986, Sadok et al. 2021, Pankasem et al. 2024). Evaporative cooling has been also tentatively proposed as the mechanism responsible for autonomous cooling in wild flowers (Herrera 2024a, Sherer et al. 2024), mainly based on the observations that flowers often bear functional stomata; similar geometric rules govern the distribution of stomata in leaves and flower parts; and transpiration through floral stomata can be greater than transpiration through leaf stomata when flowers are exposed to high temperatures (Blanke and Lovatt 1993, Azad et al. 2007, Huang et al. 2018, Zhang et al. 2018, Sinha et al. 2022). In flowers with few or nonfunctional stomata, cuticular transpiration could be another pathway for evaporation (Kitamura and Ueno 2015, Cheng et al. 2021) and contribute also to floral cooling in hot environments, since water permeability of plant cuticles increases with ambient temperature (Schönherr et al. 1979, Schönherr and Mérida 1981, Riederer and Schreiber 2001). Irrespective of the relative importance of stomata and cuticle as pathways for evaporation, the experimental results of this study showing that inert, dried capitula lacked any cooling capacity and built up a considerable thermal excess relative to the environment, provide an indication that evaporative heat exchange with the environment seems to play a central role in flower cooling just as it does in leaf cooling (Jones 1992, Lambers et al. 2008).

### Thermoregulation

The extent to which plants are able to regulate their temperatures has manifold implications in relation to environmental changes, yet “thermoregulation remains a controversial topic in plant biological research” (Drake 2023). Part of the controversy and contrasting published results could possibly be associated with the particular thermoregulatory model generally accepted.

Studies on plant thermoregulation have most often adopted to the classical definition of the concept established by Huey and Slatkin (1976) for ectothermic vertebrates, namely that thermoregulation is a process whereby a plant keeps its temperature (somewhat) independent of the temperature of the ambient. Under this general premise, a plant’s thermoregulatory capacity could be quantified inversely by the slope of the simple linear regression relating plant temperature to ambient temperature, so that slopes close to zero denote better thermoregulatory ability than slopes close to unity (Huey and Slatkin 1976). This classical linear approach has been generally deemed acceptable to depict the relationships between leaf and air temperature, and possible alternative shapes of the functional relationship between plant and ambient temperatures have been rarely explored (Campbell and Norman 1998, Michaletz et al. 2015, 2016, Miller et al. 2021, Still et al. 2022, Drake 2023, Guo, Zhang et al. 2023; but see Blonder and Michaletz 2018). Results found here for the capitula of Asteraceae, in contrast, strongly suggest that the key assumption of a single linear relationship linking plant and air temperature may not universally apply, since the relationship was best described by two linear relationships with noticeably different slopes. A formal extension of the single-slope, canonical linear model of Huey and Slatkin (1976) is introduced in Figure 7 that accomodates the two-slopes thermal models which seem to follow some flowers (present study), and possibly some leaves too (examples mentioned in the preceding section). Under this extended, two-slopes thermal regulation framework, different thermoregulatory possibilities available to plants can be formulated which differ in one or more of the following aspects: (1) the position of regression lines relative to null-model and isothermal *T*_in_–*T*_out_ relationships, both of whose slopes equal unity, (2) the comparative values of the two slopes involved, and (3) the position of break points along the gradient of environmental temperature. The species of Asteraceae studied here exemplify several of these tpossibilities (Figure 7).

Interpreting separately the two slopes obtained for each species in the light of Huey and Slatkin’s (1976) linear framework brings a similar conclusion for all species: capitula were only weakly thermoregulated (slopes close to unity) at low temperatures (*T*_out_ < Ψ), and switched to intense thermoregulation (slopes approaching zero) when ambient temperature exceeded Ψ. In some extreme cases (e.g., *Carlina*, *Carthamus*), *T*_in_ became truly independent of *T*_out_ when *T*_out_ was greater than Ψ, which denoted virtually perfect thermoregulation. In short, active floral thermoregulation was environmentally triggered when ambient temperature surpassed an species-specific thermal threshold Ψ. Declining slopes of the relationships between flower and ambient temperature as the latter increased were also reported by Rejsková et al. (2010) for *Anemone nemorosa* and *Galanthus nivalis*, two vernal flowers of temperate forest understory. This observation suggests that threshold-triggered thermoregulatory cooling would not be limited to flowers from hot environments. In addition, threshold-dependent thermoregulatory responses raise the issue of how plants sense temperature and, particularly, how they feel the heat (Franklin 2010, Ruelland and Zachowski 2010, Mittler et al. 2012), which will not be discussed here.

### Ecological and evolutionary implications

Unless plants are able to decouple intrafloral temperature from that of the surrounding air, exposure of flowers to thermal environments that are harmful or suboptimal for aspects of their reproductive function (production and viability of pollen and ovules, pollen tube growth, ovule fertilization) will eventually have a negative impact on individual fecundity (see references in Introduction). Mechanisms favoring either floral warming or floral cooling should therefore have been selected for in environments where flowers are frequently exposed, respectively, to temperatures that are either too cold or too hot for optimal performance. The former possibility has been documented in detail for many plants from cold and temperate regions (van der Kooi et al. 2019, Heiling and Koski 2024). The possibility that plants can evolve means for controlled cooling of their flowers, in contrast, has only begun to be investigated in some detail quite recently (Karban et al. 2023, Herrera 2024a, Koski et al. 2024, Sherer et al. 2024). Although this recent evidence refers to few species and environments, it supports the expectation that plants can lower floral temperature so that damage to ovules and pollen from heat stress is avoided or alleviated. Results of the present investigation represent a significant addition to this literature, for they seem the first ones from a hot environment where flowers face high temperatures for long periods on a daily basis. Thermoregulatory cooling found here was of a sufficient magnitude to have the potential to enhance reproductive performance under high ambient temperature, as estimated Ψ values were remarkably close to the average optimal temperature for pollen performance (Tushabe and Rosbakh 2024).

The frequency and duration of summer heat waves are currently increasing in the southern Iberian Peninsula (Lorenzo et al. 2021, Díaz-Poso et al. 2023, Agencia Estatal de Meteorología 2023), which highlights the value of floral cooling by the summer-flowering plants studied here. It must be stressed, however, that cooling by transpiration in these plants comes at the price of increasing water loss precisely during the harshest period of the Mediterranean dry season.

Plants could therefore face a delicate balance between undertaking sufficient short-term cooling to avoid damage to pollen and ovules, and saving sufficient water to maintain vital physiological functions during summer (Sadok et al. 2021). The increasingly warming summer climate in the study area will lead to plants spending more time at *T*_out_ > Ψ and thus increasing water expenses for capitula cooling, which could eventually reduce survival or fecundity even though their capitula had been kept within safe temperature limits most of the time. The unexplored possibility remains, however, that summer-flowering Cardueae can afford such intensive water drains because of some still unrecognized anatomical or physiological capacity to improve water acquisition, storage or retention.

Cooling of flowers can have indirect beneficial effects on both plant reproduction and pollinator populations, via enhancement of pollinator visitation arising from the appearance of a distinct, “thermally improved” microclimatic layer (Karban et al. 2023, 2024, Herrera 2024a).

The main pollinators of the plants studied here were medium- and large-sized endothermic bees in the families Megachilidae and Apidae which could risk overheating and/or dehydration while foraging in the hot-dry air of the Mediterranean summer (Corbet and Huang 2016, Herrera 2024b, Johnson et al. 2023, Kazenel et al. 2024, Rose-Person et al. 2024). Bees are able to discriminate among flowers on the basis of floral temperature, preferring to probe warmer flowers at low ambient temperature and cooler ones at ambient temperature above ∼30°C (Herrera 1995, Dyer et al. 2006, Rands and Whitney 2008, Whitney et al. 2008, Norgate et al. 2010, Harrap et al. 2017, Shrestha et al. 2018, Descamps et al. 2021). Results of the present investigation, although limited to the bee pollinators of only six plant species, are consistent with this expectation. Most bee individuals were probing capitula at ambient temperatures higher than the corresponding cooling thresholds, and the visited capitula were thus most likely to be cooler than the air. The present data, however, do not allow for rigorously proving a role of capitula temperature in itself in attracting bees, since these latter could be using floral temperature as a cue for other correlated factors such as, e.g., nectar sugar concentration (Dyer et al. 2006, Whitney et al. 2008). This caveat notwithstanding, the finding that the capitula of most locally coexisting, summer-flowering Asteraceae in my study area tended to be spatially concentrated within a narrow layer close to the ground, where air temperatures are ordinarily highest (Parry 1951, Geiger et al. 1995), highlights the potential significance at the plant community level of cooled capitula as a thermally favorable microhabitat for foraging bees during most of daytime. The possible survival and reproductive advantages derived to summer bees from foraging within that “refrigerated flower layer” remain to be investigated.

All species studied here shared the ability to internally cool their capitula, but differed in the actual descriptive thermal parameters obtained from segmented regresions, particularly the breakpoint values. The number of species examined was too small to perform a full-fledged analysis of phylogenetic signal in thermal ecology parameters, but results of variance partitions among nested taxonomic levels (Appendix S2: Table S1) provided an acceptable shortcut to it given the monophyletic nature of all the genera and tribes involved (Herrando-Moraira et al. 2019). Interspecific variance in breakpoints was taxonomically structured, with differences among subtribes being the main source of interspecific variance. This points to an early adaptive diversification of Ψ values within the Cardueae, which could perhaps have endowed this very species-rich lineage with the capacity to exploit a broad range of thermal environments. Large floral capitula are inherently prone to the accumulation of large heat loads under high ambient temperature and solar irradiance, and consequently will easily reach high, potentially harmful temperatures (see, e.g., Jones 1992 for theoretical foundations, and results of experiments with inert capitula reported here for an empirical proof). Floral thermoregulatory ability in species of Cardueae with large capitula possibly represented an adaptive breakthrough allowing the colonization of sunlit, hot-dry environments at times of year when few other plants were in bloom and competition for pollinators was probably at a yearly minimum. This speculative interpretation is supported by the fact that the earliest-diverging subtribe Carlininae (Herrando-Moraira et al. 2019) is characterized by high Ψ values and living in hotter, drier and sunnier environments than the rest of subtribes considered here. It remains to be tested whether other plant lineages displaying large flowers in hot-dry, sunny environments (e.g., Cactaceae) have also evolved facultative thermoregulatory cooling similar to that described here for the Asteraceae.

## CONCLUDING REMARKS

Plants have been long acknowledged as transformers of abiotic features of the environment in a direction that benefits other organisms (Wright and Francia 2024), or “ecosystem engineers” according to the original definition of the concept by Jones et al. (1994). As emphasized by these authors, however, separating engineering from other ecological processes may be difficult because these non-trophic interactions ordinarily co-occur with trophic interactions, a circumstance which has contributed to stir some controversies (Wright and Jones 2006). Despite discrepancies about the appropriate use of the term “ecosystem engineering”, it is currently a well-recognized type of ecological interactions (Romero et al. 2015) that can apply to the case of self-cooling, summer flower heads examined in this study. Mutualistic bees visited the cooled capitula of Asteraceae for food, as also did egg-laying females of a number of species of antagonistic seed-eating and parasitoid wasps, Tephritid flies, and Curculionid beetles (C. M. Herrera, unpublished data). Even though any microhabitat enhancement brought about by floral cooling will be linked in all these instances to trophic relationships between plants and insects, the set of flower-dwelling bees, flies, wasps and beetles could be actually benefitting from the patches of benign microclimates “engineered” by the thistles in the hot-dry summer environment. This would provide a plausible mechanism whereby plant microclimatic engineering could contribute to local insect diversity.

## ACKNOWLEDGMENTS

I am grateful to Consejería de Medio Ambiente, Junta de Andalucía, for permission to work in Sierras de Cazorla, Segura y Las Villas Natural Park and providing invaluable facilities there; Mónica Medrano for useful suggestions; and Pedro Jiménez Mejías for directions to find *Carlina gummifera*. The research reported in this paper received no specific grant from any funding agency.

## DATA AVAILABILITY STATEMENT

Data, metadata and R script will be made available at figshare

## REFERENCES

Agencia Estatal de Meteorología, AEMET. 2023. Olas de calor en España desde 1975. AEMET, Área de Climatología y Aplicaciones Operativas, Madrid. https://www.aemet.es/documentos/es/conocermas/recursos_en_linea/publicaciones_y_estudios/estudios/Olas_calor/olas_calor_actualizacionoct2023.pdf

Althawadi, A. M., and J. Grace. 1986. Water use by the desert cucurbit *Citrullus colocynthis* (L.) Schrad. Oecologia 70:475–480.

Arathi, H. S., and T. J. Smith. 2023. Drought and temperature stresses impact pollen production and autonomous selfing in a California wildflower, *Collinsia heterophylla*. Ecology and Evolution 13:e10324.

Azad, A. K., Y. Sawa, T. Ishikawa, and H. Shibata. 2007. Temperature-dependent stomatal movement in tulip petals controls water transpiration during flower opening and closing. Annals of Applied Biology 150:81–87.

Blanca, G., B. Cabezudo, M. Cueto, and C. Morales Torres, editors. 2011. Flora vascular de Andalucía oriental. 2nd edition. Universidades de Almería, Granada, Jaén y Málaga, Granada, Spain.

Blanke, M. M., and C. J. Lovatt. 1993. Anatomy and transpiration of the avocado inflorescence. Annals of Botany 71:543–547.

Blonder, B., and S. T. Michaletz. 2018. A model for leaf temperature decoupling from air temperature. Agricultural and Forest Meteorology 262:354–360.

Campbell, G. S., and J. M. Norman. 1998. An introduction to environmental biophysics. 2nd edition. Springer, New York, NY, USA.

Capel Molina, J. J. 1981. Los climas de España. Oikos-Tau, Barcelona, Spain.

Chaturvedi, P., A. J. Wiese, A. Ghatak, L. Zaveska Drabkova, W. Weckwerth, and D. Honys. 2021. Heat stress response mechanisms in pollen development. New Phytologist 231:571– 585.

Cheng, G., L. Wang, H. Wu, X. Yu, N. Zhang, X. Wan, L. He, and H. Huang. 2021. Variation in petal and leaf wax deposition affects cuticular transpiration in cut lily flowers. Frontiers in Plant Science 12:781987.

Cook, A. M., N. Berry, K. V. Milner, and A. Leigh. 2021. Water availability influences thermal safety margins for leaves. Functional Ecology 35:2179–2189.

Corbet, S. A., and S.-Q. Huang. 2016. Small bees overheat in sunlit flowers: do they make cooling flights? Ecological Entomology 41:344–350.

Descamps, C., A. Jambrek, M. Quinet, and A.-L. Jacquemart. 2021. Warm temperatures reduce flower attractiveness and bumblebee foraging. Insects 12:493.

Díaz-Poso, A., N. Lorenzo, and D. Royé. 2023. Spatio-temporal evolution of heat waves severity and expansion across the Iberian Peninsula and Balearic islands. Environmental Research 217:114864.

Drake, B. G., K. Raschke, and F. B. Salisbury. 1970. Temperature and transpiration resistances of *Xanthium* leaves as affected by air temperature, humidity, and wind speed. Plant Physiology 46:324–330.

Drake, J. E. 2023. A data-intensive documentation of plant ecosystem thermoregulation across spatial and temporal scales. New Phytologist 238:921–923.

Dyer, A. G., H. M. Whitney, S. E. J. Arnold, B. J. Glover, and L. Chittka. 2006. Bees associate warmth with floral colour. Nature 442:525–525.

Ehrler, W. L. 1973. Cotton leaf temperatures as related to soil water depletion and meteorological factors. 1. Agronomy Journal 65:404–409.

Franklin, K. A. 2010. Plant chromatin feels the heat. Cell 140:26–28.

Galen, C. 2006. Solar furnaces or swamp coolers: costs and benefits of water use by solar-tracking flowers of the alpine snow buttercup, *Ranunculus adoneus*. Oecologia 148:195– 201.

Geiger, R., R. H. Aron, and P. Todhunter. 1995. The climate near the ground. 5th ed. Vieweg, Wiesbaden, Germany.

Gómez Mercado, F. 2011. Vegetación y flora de la Sierra de Cazorla. Guineana 17:1–481.

Guo, Z., C. J. Still, C. K. F. Lee, Y. Ryu, B. Blonder, J. Wang, T. C. Bonebrake, A. Hughes, Y. Li, H. C. H. Yeung, K. Zhang, Y. K. Law, Z. Lin, and J. Wu. 2023. Does plant ecosystem thermoregulation occur? An extratropical assessment at different spatial and temporal scales. New Phytologist 238:1004–1018.

Guo, Z., K. Zhang, H. Lin, B. M. Majcher, C. K. F. Lee, C. J. Still, and J. Wu. 2023. Plant canopies exhibit stronger thermoregulation capability at the seasonal than diurnal timescales. Agricultural and Forest Meteorology 339:109582.

Harrap, M. J. M., S. A. Rands, N. Hempel de Ibarra, and H. M. Whitney. 2017. The diversity of floral temperature patterns, and their use by pollinators. eLife 6:e31262.

Heiling, J. M., and M. H. Koski. 2024. Divergent gametic thermal performance and floral warming across an elevation gradient. Evolution 78:665–678.

Hemberger, J. A., N. M. Rosenberger, and N. M. Williams. 2023. Experimental heatwaves disrupt bumblebee foraging through direct heat effects and reduced nectar production. Functional Ecology 37:591–601.

Herrando-Moraira, S., J. A. Calleja, M. Galbany-Casals, N. Garcia-Jacas, J.-Q. Liu, J. López-Alvarado, J. López-Pujol, J. R. Mandel, S. Massó, N. Montes-Moreno, C. Roquet, L. Sáez, A. Sennikov, A. Susanna, and R. Vilatersana. 2019. Nuclear and plastid DNA phylogeny of tribe Cardueae (Compositae) with Hyb-Seq data: A new subtribal classification and a temporal diversification framework. Molecular Phylogenetics and Evolution 137:313–332.

Herrera, C. M. 1995. Floral biology, microclimate, and pollination by ectothermic bees in an early-blooming herb. Ecology 76:218–228.

Herrera, C. M. 2024a. Refrigerated flowers in the torrid Mediterranean summer. Ecology 105: e4268.

Herrera, C. M. 2024b. Thermal biology diversity of bee pollinators: taxonomic, phylogenetic and plant community-level correlates. Ecological Monographs 94:e1625.

Herrera, C. M. 2024c. Thermal ecology of summer-blooming Mediterranean thistles. Dataset to be deposited at Figshare.

Herrera, C. M., A. Núñez, J. Valverde, and C. Alonso. 2023. Body mass decline in a Mediterranean community of solitary bees supports the size shrinking effect of climatic warming. Ecology 104:e4128.

Huang, X., S. Lin, S. He, X. Lin, J. Liu, R. Chen, and H. Li. 2018. Characterization of stomata on floral organs and scapes of cut ‘Real’ gerberas and their involvement in postharvest water loss. Postharvest Biology and Technology 142:39–45.

Huey, R. B., and M. Slatkin. 1976. Cost and benefits of lizard thermoregulation. Quarterly Review of Biology 51:363–384.

Johnson, M. G., K. Alvarez, and J. F. Harrison. 2023. Water loss, not overheating, limits the activity period of an endothermic Sonoran Desert bee. Functional Ecology 37:2855–2867.

Jones, C. G., J. H. Lawton, and M. Shachak. 1994. Organisms as ecosystem engineers. Oikos 69:373–386.

Jones, H. G. 1992. Plants and microclimate. 2nd edition. Cambridge University Press, Cambridge, UK.

Karban, R., D. Rutkowski, and N. A. Murray. 2023. Flowers that self-shade reduce heat stress and pollen limitation. American Journal of Botany 110:e16109.

Karban, R., M. Huntzinger, D. Rutkowski, and N. Murray. 2024. Petal-shading in *Romneya coulteri* affects seed set and interactions with floral visitors. Arthropod-Plant Interactions 18:1065–1073.

Kazenel, M. R., K. W. Wright, T. Griswold, K. D. Whitney, and J. A. Rudgers. 2024. Heat and desiccation tolerances predict bee abundance under climate change. Nature 628:342–348.

Kitamura, Y., and S. Ueno. 2015. Inhibition of transpiration from the inflorescence extends the vase life of cut Hydrangea flowers. Horticulture Journal 84:156–160.

Koski, M. H., J. M. Heiling, and J. S. Apland. 2024. Behavioural thermoregulation of flowers via petal movement. Ecology Letters 27:e14524.

Lambers, H., F. S. Chapin, and T. L. Pons. 2008. Plant physiological ecology. 2nd edition. Springer, New York, NY, USA.

Linacre, E. T. 1964. A note on a feature of leaf and air temperatures. Agricultural Meteorology 1:66–72.

Linacre, E. T. 1967. Further notes on a feature of leaf and air temperatures. Archiv für Meteorologie, Geophysik und Bioklimatologie, Serie B 15:422–436.

Lohani, N., M. B. Singh, and P. L. Bhalla. 2020. High temperature susceptibility of sexual reproduction in crop plants. Journal of Experimental Botany 71:555–568.

Lorenzo, N., A. Díaz-Poso, and D. Royé. 2021. Heatwave intensity on the Iberian Peninsula: future climate projections. Atmospheric Research 258:105655.

Manzi, O. J. L., M. Wittemann, M. E. Dusenge, J. Habimana, A. Manishimwe, M. Mujawamariya, B. Ntirugulirwa, E. Zibera, L. Tarvainen, D. Nsabimana, G. Wallin, and J. Uddling. 2024. Canopy temperatures strongly overestimate leaf thermal safety margins of tropical trees. New Phytologist 243:2115–2129.

Michaletz, S. T., M. D. Weiser, J. Zhou, M. Kaspari, B. R. Helliker, and B. J. Enquist. 2015. Plant thermoregulation: energetics, trait–environment interactions, and carbon economics. Trends in Ecology & Evolution 30:714–724.

Michaletz, S. T., M. D. Weiser, N. G. McDowell, J. Zhou, M. Kaspari, B. R. Helliker, and B. J. Enquist. 2016. The energetic and carbon economic origins of leaf thermoregulation. Nature Plants 2:16129.

Miller, B. D., K. R. Carter, S. C. Reed, T. E. Wood, and M. A. Cavaleri. 2021. Only sun-lit leaves of the uppermost canopy exceed both air temperature and photosynthetic thermal optima in a wet tropical forest. Agricultural and Forest Meteorology 301–302:108347.

Mittler, R., A. Finka, and P. Goloubinoff. 2012. How do plants feel the heat? Trends in Biochemical Sciences 37:118–125.

Muggeo, V. M. R. 2008. segmented: an R package to fit regression models with broken-line relationships. R News 8: 20–25.

Muggeo, V. M. R. 2016. Testing with a nuisance parameter present only under the alternative: a score-based approach with application to segmented modelling. Journal of Statistical Computation and Simulation 86:3059–3067.

Muggeo, V. M. R. 2017. Interval estimation for the breakpoint in segmented regression: a smoothed score-based approach. Australian & New Zealand Journal of Statistics 59:311– 322.

Norgate, M., S. Boyd-Gerny, V. Simonov, M. G. P. Rosa, T. A. Heard, and A. G. Dyer. 2010. Ambient temperature influences Australian native stingless bee (*Trigona carbonaria*) preference for warm nectar. PLoS ONE 5:e12000.

Palmer, J. H. 1967. Diurnal variation in leaf and boll temperatures of irrigated cotton grown under two soil moisture regimes in a semi-arid climate. Agricultural Meteorology 4:39–54.

Pankasem, N., P.-K. Hsu, B. N. K. Lopez, P. J. Franks, and J. I. Schroeder. 2024. Warming triggers stomatal opening by enhancement of photosynthesis and ensuing guard cell CO sensing, whereas higher temperatures induce a photosynthesis-uncoupled response. New Phytologist 244:1847–1863.

Parry, D. A. 1951. Microclimate close to the ground. Nature 167:73–74.

Patiño, S., and J. Grace. 2002. The cooling of convolvulaceous flowers in a tropical environment. Plant, Cell & Environment 25:41–51.

Pearcy, R. W., J. A. Berry, and B. Bartholomew. 1972. Field measurements of the gas exchange capacities of *Phragmites communis* under summer conditions in Death Valley. Carnegie Institution Year Book 71:161–164.

Perkins-Kirkpatrick, S. E., and S. C. Lewis. 2020. Increasing trends in regional heatwaves. Nature Communications 11:3357.

Pinheiro, J., D. Bates, and R Core Team. 2024. nlme: linear and nonlinear mixed effects models. R package version 3.1–166, https://CRAN.R-project.org/package=nlme.

Posch, B. C., S. E. Bush, D. F. Koepke, A. Schuessler, L. L. D. Anderegg, L. M. T. Aparecido, B. W. Blonder, J. S. Guo, K. L. Kerr, M. E. Moran, H. F. Cooper, C. E. Doughty, C. A. Gehring, T. G. Whitham, G. J. Allan, and K. R. Hultine. 2024. Intensive leaf cooling promotes tree survival during a record heatwave. Proceedings of the National Academy of Sciences USA 121:e2408583121.

Pugnaire, F. I., M. Arista, J. S. Carrión, J. A. Devesa, C. M. Herrera, G. Nieto Feliner, P. J. Rey, and C. Alonso. 2024. Origen y retos actuales de la diversidad vegetal en las sierras Béticas, un área de diversidad de importancia global. Ecosistemas 33:2676.

R Core Team. 2024. R: A language and environment for statistical computing. Vienna, Austria: R Foundation for Statistical Computing.

Rands, S. A., and H. M. Whitney. 2008. Floral temperature and optimal foraging: is heat a feasible floral reward for pollinators? PLoS ONE 3:2007.

Rejsková, A., J. Brom, J. Pokorny, and J. Korecko. 2010. Temperature distribution in light-coloured flowers and inflorescences of early spring temperate species measured by infrared camera. Flora 205:282–289.

Riederer, M., and L. Schreiber. 2001. Protecting against water loss: analysis of the barrier properties of plant cuticles. Journal of Experimental Botany 52:2023–2032.

Romero, G. Q., T. Gonçalves Souza, C. Vieira, and J. Koricheva. 2015. Ecosystem engineering effects on species diversity across ecosystems: a meta analysis. Biological Reviews 90:877– 890.

Rosbakh, S., E. Pacini, M. Nepi, and P. Poschlod. 2018. An unexplored side of regeneration niche: seed quantity and quality are determined by the effect of temperature on pollen performance. Frontiers in Plant Science 9:1036.

Rose-Person, A., L. S. Santiago, and N. E. Rafferty. 2024. Drought stress influences foraging preference of a solitary bee on two wildflowers. Annals of Botany. Doi:10.1093/aob/mcae048

Rosenberger, N. M., J. A. Hemberger, and N. M. Williams. 2024. Heatwaves exacerbate pollen limitation through reductions in pollen production and pollen vigour. AoB Plants 16:plae045.

Ruelland, E., and A. Zachowski. 2010. How plants sense temperature. Environmental and Experimental Botany 69:225–232.

Ryan, S. E., and L. S. Porth. 2007. A tutorial on the piecewise regression approach applied to bedload transport data. General Technical Report RMRS-GTR-189. Fort Collins, CO: U.S. Department of Agriculture, Forest Service, Rocky Mountain Research Station. 41 p.

Sadok, W., J. R. Lopez, and K. P. Smith. 2021. Transpiration increases under high temperature stress: potential mechanisms, trade offs and prospects for crop resilience in a warming world. Plant, Cell & Environment 44:2102–2116.

Schönherr, J., K. Eckl, and H. Gruler. 1979. Water permeability of plant cuticles: the effect of temperature on diffusion of water. Planta 147:21–26.

Schönherr, J., and T. Mérida. 1981. Water permeability of plant cuticular membranes: the effects of humidity and temperature on the permeability of non isolated cuticles of onion bulb scales. Plant, Cell & Environment 4:349–354.

Seymour, R. S., Y. Ito, Y. Onda, and K. Ito. 2009. Effects of floral thermogenesis on pollen function in Asian skunk cabbage *Symplocarpus renifolius*. Biology Letters 5:568–570.

Sherer, T. N., J. M. Heiling, and M. H. Koski. 2024. Floral thermal biology in relation to pollen thermal performance in an early spring flowering plant. Plant Biology 26:811–820.

Shrestha, M., J. E. Garcia, Z. Bukovac, A. Dorin, and A. G. Dyer. 2018. Pollination in a new climate: assessing the potential influence of flower temperature variation on insect pollinator behaviour. PLoS ONE 13:e0200549.

Sinha, R., S. I. Zandalinas, Y. Fichman, S. Sen, S. Zeng, A. Gómez Cadenas, T. Joshi, F. B. Fritschi, and R. Mittler. 2022. Differential regulation of flower transpiration during abiotic stress in annual plants. New Phytologist 235:611–629.

Skubatz, H. 2014. Thermoregulation in the appendix of the *Sauromatum guttatum* (Schott) inflorescence. Botanical Studies 55:1–13.

Smith, W. K. 1978. Temperatures of desert plants: another perspective on the adaptability of leaf size. Science 201:614–616.

Still, C. J., G. Page, B. Rastogi, D. M. Griffith, D. M. Aubrecht, Y. Kim, S. P. Burns, C. V. Hanson, H. Kwon, L. Hawkins, F. C. Meinzer, S. Sevanto, D. Roberts, M. Goulden, S. Pau, M. Detto, B. Helliker, and A. D. Richardson. 2022. No evidence of canopy-scale leaf thermoregulation to cool leaves below air temperature across a range of forest ecosystems. Proceedings of the National Academy of Sciences USA 119:e2205682119.

Tarvainen, L., M. Wittemann, M. Mujawamariya, A. Manishimwe, E. Zibera, B. Ntirugulirwa, C. Ract, O. J. L. Manzi, M. X. Andersson, C. Spetea, D. Nsabimana, G. Wallin, and J. Uddling. 2022. Handling the heat – photosynthetic thermal stress in tropical trees. New Phytologist 233:236–250.

Toms, J. D., and M. L. Lesperance. 2003. Piecewise regression: a tool for identifying ecological thresholds. Ecology 84:2034–2041.

Tushabe, D., and S. Rosbakh. 2024. Patterns and drivers of pollen temperature tolerance. Plant, Cell & Environment. Doi:10.1111/pce.15207

Upchurch, D. R., and J. R. Mahan. 1988. Maintenance of constant leaf temperature by plants. II. Experimental observations in cotton. Environmental and Experimental Botany 28:359–366.

van der Kooi, C. J., P. G. Kevan, and M. H. Koski. 2019. The thermal ecology of flowers. Annals of Botany 124:343–353.

Whitney, H. M., A. Dyer, L. Chittka, S. A. Rands, and B. J. Glover. 2008. The interaction of temperature and sucrose concentration on foraging preferences in bumblebees. Naturwissenschaften 95:845–850.

Wright, A. J., and R. M. Francia. 2024. Plant traits, microclimate temperature and humidity: A research agenda for advancing nature-based solutions to a warming and drying climate. Journal of Ecology 112:2462–2470.

Wright, J. P., and C. G. Jones. 2006. The concept of organisms as ecosystem engineers ten years on: progress, limitations, and challenges. BioScience 56:203–209.

Zhang, F. P., M. R. Carins Murphy, A. A. Cardoso, G. J. Jordan, and T. J. Brodribb. 2018. Similar geometric rules govern the distribution of veins and stomata in petals, sepals and leaves. New Phytologist 219:1224–1234.

