## Appendix S1 for "Flowers that cool themselves: thermal ecology of summer-blooming thistles in hot Mediterranean environments"

**Appendix S1. Supplementary material to the section Materials and Methods**

**Figure S1.** Examples of thermocouple setup for the continuous measurement of paired *T*_in_–*T*_out_ temperatures.

| 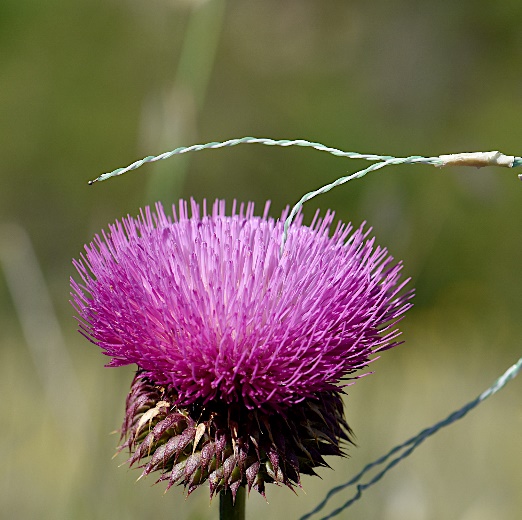  *Carduus granatensis* | 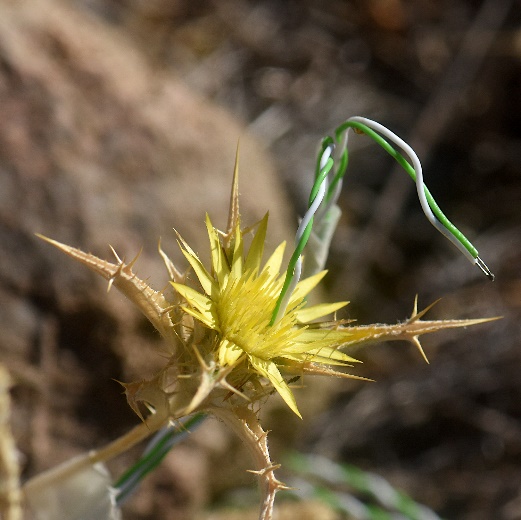  *Carlina corymbosa* |
| --- | --- |
| 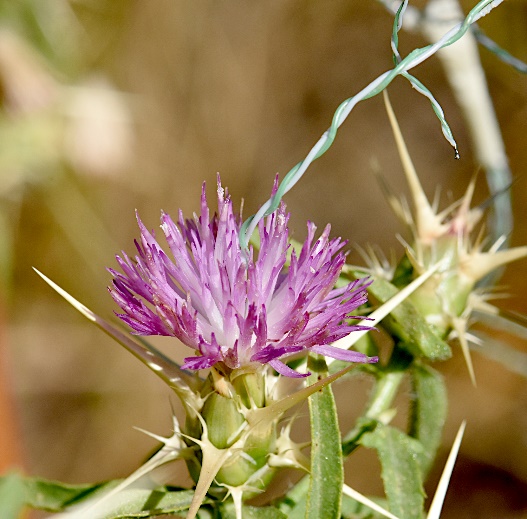  *Centaurea calcitrapa* | 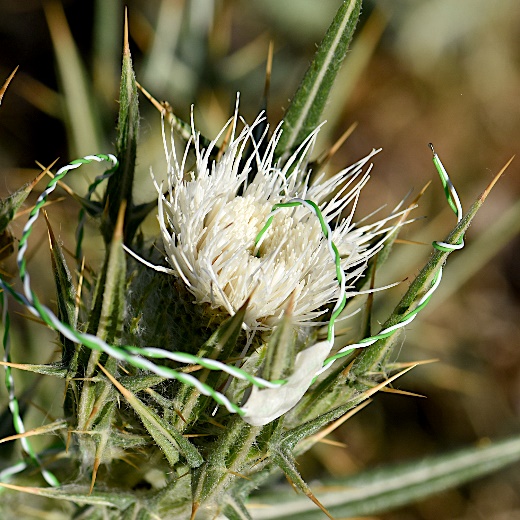  *Cirsium odontolepis* |
| 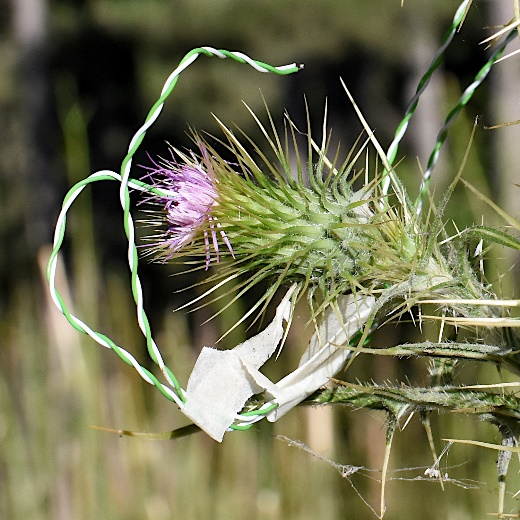  *Cirsium vulgare* | 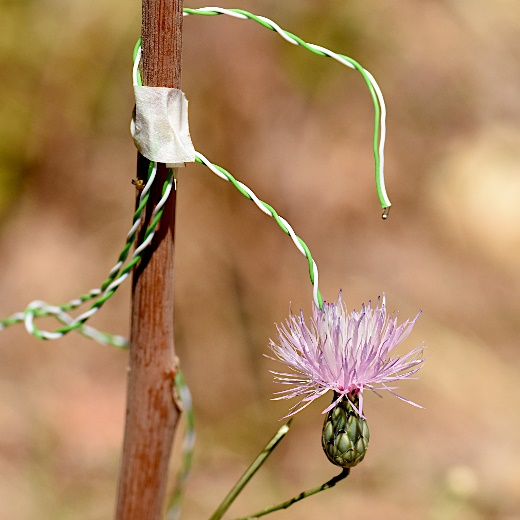  *Mantisalca salmantica* |

**Figure S2.** Fast-dried floral capitula arranged for measurements of paired *T*_in_–*T*_out_ temperatures in the field.

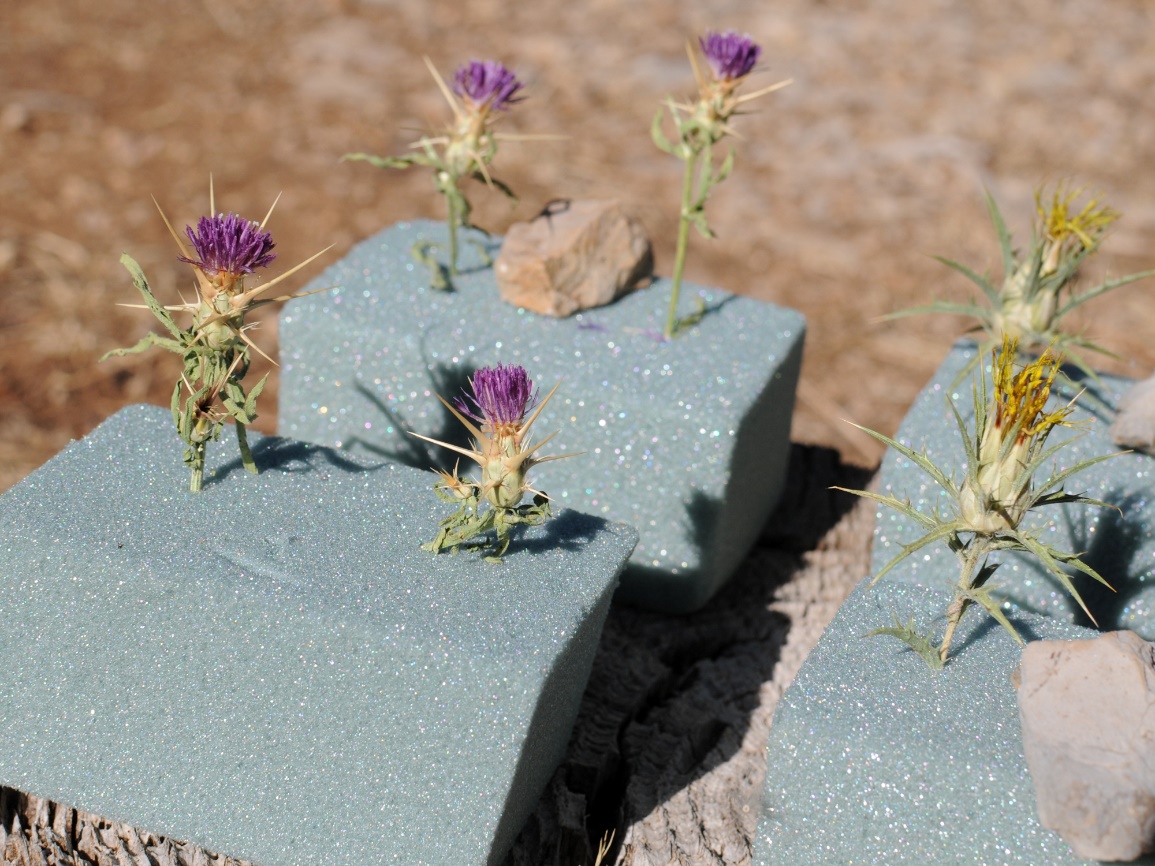

*Centaurea calcitrapa* (left) and *Carthamus lanatus* (right)

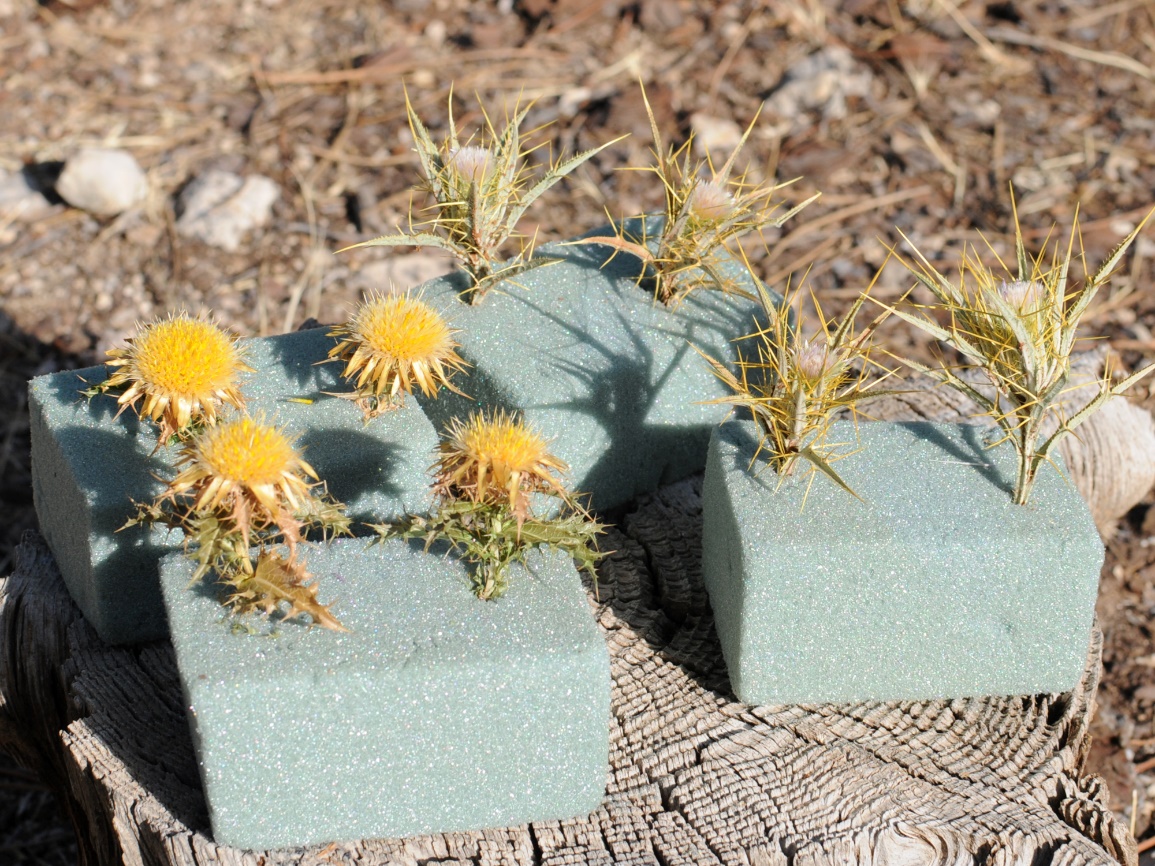

*Carlina hispanica* (left) and *Picnomon acarna* (right)

**Table S1.** Type of data collected for each of the 15 species considered in the study.

|  | Data type | | | | |
| --- | --- | --- | --- | --- | --- |
| Species | Experimental capitula | Instantaneous measurements | Continuous monitoring | Vertical distribution | Bee visitation |
| *Carduus granatensis* |  | **✓** | **✓** |  |  |
| *Carlina gummifera* |  | **✓** |  | **✓** |  |
| *Carlina hispanica* | **✓** | **✓** | **✓** | **✓** | **✓** |
| *Carlina racemosa* | **✓** | **✓** | **✓** | **✓** |  |
| *Carlina vulgaris* |  |  | **✓** |  |  |
| *Carthamus lanatus* | **✓** | **✓** |  | **✓** |  |
| *Centaurea calcitrapa* | **✓** | **✓** | **✓** | **✓** | **✓** |
| *Centaurea ornata* |  | **✓** |  |  |  |
| *Cirsium acaule* |  | **✓** | **✓** | **✓** |  |
| *Cirsium odontolepis* |  | **✓** | **✓** |  | **✓** |
| *Cirsium pyrenaicum* | **✓** | **✓** | **✓** | **✓** | **✓** |
| *Cirsium vulgare* | **✓** | **✓** | **✓** | **✓** | **✓** |
| *Mantisalca salmantica* | **✓** | **✓** | **✓** | **✓** | **✓** |
| *Picnomon acarna* | **✓** | **✓** | **✓** | **✓** |  |
| *Ptilostemon hispanicus* | **✓** | **✓** | **✓** | **✓** |  |
