## Appendix S2 for "Flowers that cool themselves: thermal ecology of summer-blooming thistles in hot Mediterranean environments"

**Appendix S2. Supplementary material to the Results section.**

Table S1. Partition of variance among taxonomic levels of the interspecific variation in thermal parameters from segmented regressions of *T*_in_ against *T*_out_ (Table 1).

|  | Variance component | | |
| --- | --- | --- | --- |
| Segmented regression parameter | Among subtribes | Genera nested within subtribes | Species nested within genera |
| Ψ | 8.52 | 0.80 | 2.63 |
| Slope 1 | 0 | 0 | 0.048 |
| Slope 2 | 0 | 0.012 | 0.009 |

Table S2. Bees observed foraging on the capitula of six species of Asteraceae, broken down by bee and plant species (*N* = number of individuals). Air temperature at the foraging site was measured for each individual bee.

| Plant species | Bee species | *N* |
| --- | --- | --- |
| *Carlina hispanica* | *Amegilla albigena* | 6 |
| *Carlina hispanica* | *Amegilla quadrifasciata* | 7 |
| *Carlina hispanica* | *Ammobates punctatus* | 2 |
| *Carlina hispanica* | *Anthophora fulvodimidiata* | 35 |
| *Carlina hispanica* | *Halictus fulvipes* | 12 |
| *Carlina hispanica* | *Halictus pollinosus* | 3 |
| *Carlina hispanica* | *Halictus scabiosae* | 5 |
| *Carlina hispanica* | *Halictus subauratus* | 11 |
| *Carlina hispanica* | *Hylaeus variegatus* | 1 |
| *Carlina hispanica* | *Icteranthidium laterale* | 6 |
| *Carlina hispanica* | *Lithurgus chrysurus* | 1 |
| *Carlina hispanica* | *Megachile albisecta* | 5 |
| *Carlina hispanica* | *Megachile apicalis* | 2 |
| *Carlina hispanica* | *Megachile lagopoda* | 1 |
| *Carlina hispanica* | *Megachile octosignata* | 1 |
| *Carlina hispanica* | *Megachile pilidens* | 3 |
| *Carlina hispanica* | *Megachile pilicrus* | 1 |
| *Carlina hispanica* | *Nomiapis paulyi* | 1 |
| *Carlina hispanica* | *Pseudoanthidium scapulare* | 4 |
| *Carlina hispanica* | *Xylocopa cantabrita* | 1 |
| *Carlina hispanica* | *Xylocopa violacea* | 3 |
| *Centaurea calcitrapa* | *Amegilla albigena* | 21 |
| *Centaurea calcitrapa* | *Amegilla ochroleuca* | 1 |
| *Centaurea calcitrapa* | *Amegilla quadrifasciata* | 32 |
| *Centaurea calcitrapa* | *Ammobates punctatus* | 1 |
| *Centaurea calcitrapa* | *Bombus terrestris* | 1 |
| *Centaurea calcitrapa* | *Heriades crenulatus* | 6 |
| *Centaurea calcitrapa* | *Icteranthidium grohmanni* | 1 |
| *Centaurea calcitrapa* | *Lithurgus chrysurus* | 69 |
| *Centaurea calcitrapa* | *Megachile albisecta* | 56 |
| *Centaurea calcitrapa* | *Megachile apicalis* | 20 |
| *Centaurea calcitrapa* | *Megachile melanopyga* | 7 |
| *Centaurea calcitrapa* | *Megachile octosignata* | 1 |
| *Centaurea calcitrapa* | *Megachile pilicrus* | 5 |
| *Centaurea calcitrapa* | *Pseudoanthidium melanurum* | 3 |
| *Centaurea calcitrapa* | *Tetraloniella julliani* | 2 |
| *Centaurea calcitrapa* | *Thyreus histrionicus* | 1 |
| *Cirsium odontolepis* | *Halictus scabiosae* | 1 |
| *Cirsium pyrenaicum* | *Anthidium taeniatum* | 1 |
| *Cirsium pyrenaicum* | *Coelioxys afer* | 1 |
| *Cirsium pyrenaicum* | *Epeolus fallax* | 13 |
| *Cirsium pyrenaicum* | *Halictus scabiosae* | 3 |
| *Cirsium pyrenaicum* | *Icteranthidium laterale* | 16 |
| *Cirsium pyrenaicum* | *Megachile octosignata* | 6 |
| *Cirsium pyrenaicum* | *Pseudoanthidium scapulare* | 2 |
| *Cirsium vulgare* | *Amegilla quadrifasciata* | 2 |
| *Cirsium vulgare* | *Icteranthidium laterale* | 1 |
| *Cirsium vulgare* | *Megachile albisecta* | 1 |
| *Cirsium vulgare* | *Pseudoanthidium melanurum* | 1 |
| *Cirsium vulgare* | *Xylocopa violacea* | 1 |
| *Mantisalca salmantica* | *Megachile centuncularis* | 1 |
